# Clonal replacement of tumor-specific T cells following PD-1 blockade

**DOI:** 10.1101/648899

**Authors:** Kathryn E. Yost, Ansuman T. Satpathy, Daniel K. Wells, Yanyan Qi, Chunlin Wang, Robin Kageyama, Katherine McNamara, Jeffrey M. Granja, Kavita Y. Sarin, Ryanne A. Brown, Rohit K. Gupta, Christina Curtis, Samantha L. Bucktrout, Mark M. Davis, Anne Lynn S. Chang, Howard Y. Chang

## Abstract

Immunotherapies that block inhibitory checkpoint receptors on T cells have transformed the clinical care of cancer patients. However, which tumor-specific T cells are mobilized following checkpoint blockade remains unclear. Here, we performed paired single-cell RNA- and T cell receptor (TCR)-sequencing on 79,046 cells from site-matched tumors from patients with basal cell carcinoma (BCC) or squamous cell carcinoma (SCC) pre- and post-anti-PD-1 therapy. Tracking TCR clones and transcriptional phenotypes revealed a coupling of tumor-recognition, clonal expansion, and T cell dysfunction: the T cell response to treatment was accompanied by clonal expansions of CD8^+^CD39^+^ T cells, which co-expressed markers of chronic T cell activation and exhaustion. However, this expansion did not derive from pre-existing tumor infiltrating T cell clones; rather, it comprised novel clonotypes, which were not previously observed in the same tumor. Clonal replacement of T cells was preferentially observed in exhausted CD8^+^ T cells, compared to other distinct T cell phenotypes, and was evident in BCC and SCC patients. These results, enabled by single-cell multi-omic profiling of clinical samples, demonstrate that pre-existing tumor-specific T cells may be limited in their capacity for re-invigoration, and that the T cell response to checkpoint blockade relies on the expansion of a distinct repertoire of T cell clones that may have just recently entered the tumor.

## Introduction

Clinical therapies that enhance the activity of tumor-specific T cells have demonstrated efficacy in a variety of human cancers^1^. However, these immunotherapies are not effective in all patients with a given cancer subtype, nor in all cancers^2, 3^. For example, while a subset of patients with metastatic melanoma experience a response to immunotherapy based on blocking the T cell inhibitory receptor programmed death-1 (PD-1), including clinical remission up to 10 years after treatment, nearly 70% of patients do not respond or relapse shortly after remission^4–6^. Therefore, studies interrogating the molecular basis for successful clinical and immunological responses to therapy are needed.

The efficacy of the anti-tumor T cell response relies on several mechanisms, including the presentation of tumor antigens by major histocompatibility complex (MHC) molecules, clonal expansion of tumor-specific T cells, and the sustained cytolytic activity of these cells for the duration of treatment. Deficiencies in any of these steps can lead to an ineffective or incomplete response and cancer progression. For example, tumor-infiltrating T lymphocytes (TILs) can undergo ‘T cell exhaustion,’ a differentiation process that induces the upregulation of inhibitory surface receptors, decreases activation in response to cognate antigens, and limits the potential for reinvigoration following PD-1 blockade^7–12^. However, since T cell clones bearing tumor-specific T cell receptors (TCR) represent only a minority of all TILs^13^, the measurement of such dysfunction-associated molecular pathways can be challenging in ensemble cell profiling experiments. Moreover, whether dysfunctional states limit the reinvigoration of tumor-intrinsic T cells by checkpoint blockade, and whether successful immunotherapy relies on overcoming such dysfunction remains unclear.

These observations underscore the importance of single-cell analysis for dissecting the clonal T cell response to cancer. However, to date, such studies have not been adequately performed in site-matched human tumor samples following immunotherapy response, partly due the difficulty in obtaining the necessary clinical specimens. Therefore, the molecular profiles of TILs following immunotherapy and their clonal dynamics pre- and post-treatment remain poorly understood. Here we describe the single-cell landscape of advanced basal cell carcinoma (BCC) and squamous cell carcinoma (SCC) before and after anti-PD-1 treatment in site-matched primary tumors. We integrate scRNA- and scTCR-seq methods to show several principles of the clonal T cell response to checkpoint blockade in these tumor types: 1) T cells expressing the identical TCR exhibit highly correlated phenotypes, which are maintained after PD-1 blockade, 2) clonally-expanded T cells are enriched in CD8^+^CD39^+^ cells post-therapy and co-express markers of chronic T cell activation and exhaustion, and 3) major pre-therapy T cell clones are not reinvigorated following PD-1 blockade, but rather are replaced by novel expanded TCR clones that were not previously observed in the same tumor. These results suggest that checkpoint blockade relies on the recruitment of tumor-extrinsic T cell clones, which has implications for response monitoring and prediction, as well as for the future design of immunotherapies based on the premise of recruiting new waves of T cell clones to the tumor microenvironment (TME).

## Results

We sampled site-matched BCCs pre- and post-PD-1 blockade from 11 patients, many of whom were enrolled in a proof-of-principle, non-randomized, open-label study of pembrolizumab (anti-PD-1)^14^, and generated droplet-based 5’ scRNA- and TCR-seq libraries (24 samples total) (**Fig. 1a, Supplemental Table 1, and Methods**). All patients had histologically-proven advanced or metastatic BCC and were not good candidates for surgical resection^14^. Exclusion criteria included prior exposure to checkpoint blockade agents, use of systemic immunosuppressants within 4 weeks of first biopsy, or treatment with radiotherapy or any other anti-cancer treatment within 4 weeks of first biopsy^14^. Pretherapy biopsies were collected from accessible skin tumors either on Day 0 (prior to anti-PD-1 blockade initiation) or during prior visits. Post-therapy specimens were collected from the same tumor site an average of ~2 months after treatment, and best response was assessed using RECIST criteria^15^ with 55% of patients demonstrating a clinical response, comparable with published results^14^ (**Supplementary Table 1**).

**Figure 1.**
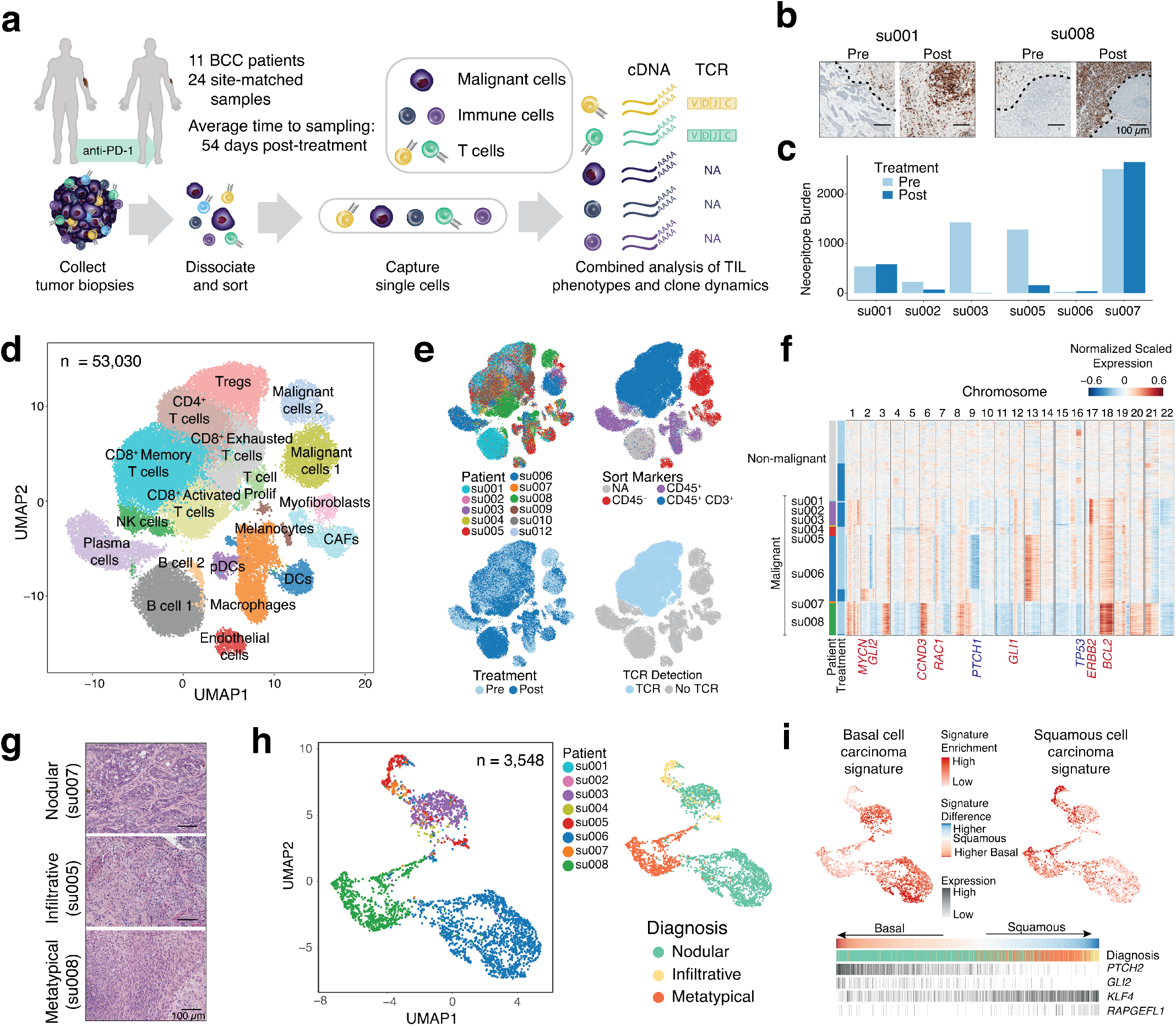
Characterization of the BCC TME pre- and post-PD-1 blockade by singlecell RNA-seq. **(a)** Workflow for sample processing and scRNA-seq analysis of advanced BCC samples collected pre- and post-PD-1 blockade. **(b)** Immunohistochemistry staining for CD3^+^ cells in representative BCC tumors before and after PD-1 blockade. Tumor boundaries denoted with dashed lines. All scale bars represent 100 μm. **(c)** Bar plot of neoepitope burden pre- and post-treatment based on exome sequencing. Variants were classified as predicted neoepitopes if the peptide was found to bind to the MHC allele with less than 500 nM binding strength and its wildtype cognate bound to the same allele with greater than 500 nM binding strength. **(d)** UMAP of all tumor-resident cells pre- and posttherapy for all 11 BCC patients. Clusters denoted by color were identified by shared nearest neighbor (SNN) clustering and labeled with inferred cell types. CAFs: cancer-associated fibroblasts, DCs: dendritic cells, pDCs: plasmacytoid dendritic cells. **(e)** UMAP of tumor-resident cells colored by patient identity (top left), FACS sort markers (top right), anti-PD1 treatment status (bottom left), and TCR detection (bottom right). **f)** Inferred CNV profiles based on scRNA-seq data. Non-immune, non-malignant cells (fibroblasts and endothelial cells, n = 2,122) were used as normal reference for malignant cell CNV inference (n = 3,548). **(g)** Representative examples of hematoxylin and eosin (H&E) staining of different BCC subtypes. All scale bars represent 100 μm. **(h)** UMAP of malignant cells colored by patient (left) and clinical subtype (right). **(i)** UMAP of malignant cells colored by enrichment of basal and squamous cell carcinoma gene signatures (from Atwood *et al*., 2015 and Hoang *et al*., 2017) (top). Malignant cells ordered based on the difference between basal and squamous signatures (bottom). The clinical diagnosis associated with each cell and expression of signature associated genes are indicated below.

We also performed several orthogonal measurements, including H&E staining and immunohistochemistry (IHC) for CD3, CD8, and PDL1, whole exome sequencing (WES), HLA typing, and bulk tumor and peripheral blood TCR-seq for the majority of samples (**Fig. 1b-c and Supplementary Table 1**). These data supported the presence of an immunological response to checkpoint blockade. First, histological assessments showed a diffuse stromal distribution of CD3^+^ T cells prior to checkpoint blockade and increased T cell infiltration following therapy (**Fig. 1b**). Second, WES revealed a high mutational burden characteristic of BCC (average 17 mutations/Mb, range 0.25-58 mutations/Mb), as well as mutations in known driver genes^16^ and a strong bias for C to T mutations characteristic of UV-induced mutations in skin cancers (**Supplementary Fig. 1a**). Analysis of mutation burden pre- and post-treatment from patients with matched WES identified several patients with significant mutational loss following treatment affecting both clonal and sub-clonal mutations, as well as neoepitopes, suggesting the presence of tumor immunoediting following PD-1 blockade (**Fig. 1c, Supplementary Fig. 1b-c**)^17^. Interestingly, we observed two cases of novel tumor sub-clones emerging post-treatment which were devoid of predicted neoepitopes (**Supplementary Fig. 1c**).

In total, we obtained scRNA-seq profiles from 55,666 malignant, immune, and stromal cells from BCC samples, of which 53,030 cells passed quality control filters (95%; **Fig. 1d, Supplementary Fig. 1d and Methods**). We performed a second cDNA amplification step to amplify TCR sequences and obtained *TRA* or *TRB* reads from 28,371/33,106 T cells (85%), of which 19,296 cells (68%) generated paired *TRA* and *TRB* data (**Fig. 1e and Methods**). We first broadly clustered single cells by scRNA-seq profiles using shared nearest neighbor (SNN) clustering based on significant principal components (PCs) and visualized cell clusters using UMAP projection (**Fig. 1d, Methods**)^18–20^. This method identified 19 clusters, which were annotated as 2 malignant cell clusters, 6 T cell clusters (2 CD4^+^ T cell clusters, 3 CD8^+^ T cell clusters, and proliferating T cells), 4 stromal cell clusters (endothelial cells, melanocytes, myofibroblasts, and cancer-associated fibroblasts), 3 myeloid clusters (dendritic cells, macrophages, and plasmacytoid dendritic cells), 3 B cell clusters (2 B cell clusters and plasma cells), and 1 NK cell cluster (**Fig. 1d**). Cell types were annotated using three parallel approaches: 1) correlation between aggregated cluster expression profiles and bulk RNA expression profiles from reference populations^21^, 2) differential gene expression, and 3) known marker gene expression (**Methods, Supplementary Fig. 2a-c**). Immune cell classifications based on gene expression profiles were consistent with the staining characteristics of labeled antibodies to cell surface markers used to isolate cells prior to scRNA-seq, and TCR sequences were almost exclusively detected in cells classified as T cells (**Fig. 1e, Supplementary Fig. 2d**). Notably, immune cells from different patients clustered together, indicating that immune cell types in the BCC tumor microenvironment (TME) were largely consistent across patients and did not represent patient-specific subpopulations or batch effects (**Fig. 1e, Supplementary Fig. 2d**). We confirmed malignant cell classification using single-cell copy number variation (CNV) estimation relative to non-malignant stromal cells (**Methods**)^22^. This analysis revealed patient-specific copy alterations only in malignant cell clusters, which were consistent with CNVs detected by bulk tumor exome sequencing, as well as with previously described CNVs in BCC (**Fig. 1f, Supplementary Fig. 3a**)^16^.

The samples in this study included the full spectrum of BCC subtypes, including nodular, infiltrative, and metatypical subtypes, which are characterized by differences in histopathology and clinical outcome (**Fig. 1g, Supplementary Table 1**)^23^. To explore gene expression associated with BCC subtypes, we re-clustered 3,548 malignant cells and found that malignant cells segregated by patient and by subtype (**Fig. 1h**), indicating significant intertumoral heterogeneity, as has been observed in other cancer types^24, 25^. We identified a core tumor gene expression program that was consistent across tumor cells from all patients and distinguished malignant cells from non-malignant cells based on differential expression (**Supplementary Fig. 3b**). This gene signature included *EPCAM, BCAM* and *TP63*, which are diagnostic markers of BCC and encode key regulators of cellular adhesion and epithelial identity (**Supplementary Fig. 3b**)^26–28^. We also identified 577 genes differentially expressed across patient tumors, including genes related to Ras signaling, suggesting aberrant activation of squamous cell pathways, as has been suggested previously (**Supplementary Fig. 3b**)^29^. To explore this further, we scored individual malignant cells for enrichment of BCC and squamous cell carcinoma (SCC) gene expression signatures derived from bulk data^30, 31^. This analysis revealed mutually exclusive gene expression patterns in individual tumors (**Fig. 1i**). Namely, tumors cells from individual patients existed along a spectrum from basal cell to squamous cell differentiation. Ordering of cells based on signature scores revealed an association of differentiation signatures and histological presentation; nodular BCCs were enriched for the basal cell signature, while infiltrative and metatypical BCCs were enriched for the squamous signature (**Fig. 1i**). Altogether, these results demonstrate that BCC gene expression is driven by patient-specific malignant pathways, but largely does not influence immune cell infiltration patterns in the TME.

We next focused our analysis on TILs to understand the clonal T cell response to checkpoint blockade using paired scRNA and TCR-seq (scRNA_+_TCR-seq). First, we reclustered 33,106 TILs and identified 9 distinct T cell clusters (**Fig. 2a, Supplementary Fig. 4a-c**). These included 3 predominantly CD4^+^ T cell clusters, 4 predominantly CD8^+^ T cell clusters, and 2 clusters that included both subtypes. All clusters contained cells from multiple patients and both pre- and post-treatment timepoints (**Fig. 2a, Supplementary Fig. 4d**). We annotated cluster phenotypes using differentially expressed marker genes and comparisons to bulk RNA-seq data^13^ (**Methods, Fig. 2b, Supplementary Fig. 4a-c**). CD4^+^ clusters included: 1) regulatory T cells (Tregs), which expressed *FOXP3, IL2RA*, and *CTLA4*, 2) follicular helper T cells (Tfh), which expressed *BTLA* and *CD200*, and 3) T helper 17 cells (Th17), which expressed *IL26* and *KLRB1* (**Fig. 2a-b**). CD8^+^ T cell clusters spanned the spectrum of T cell activation and included: 1) naïve cells, which expressed *CCR7* and *IL7R*, 2) memory cells, which expressed *EOMES* and *CXCR3*, 3) effector memory cells, which expressed memory cell genes and effector genes (*FGFBP2* and *KLRD1*), 4) activated cells, which expressed cytotoxic genes (*IFNG* and *TNF*) and AP-1 transcription factors (*FOS, JUN*), and 5) chronically activated/exhausted cells (hereafter referred to as exhausted cells), which expressed known markers of T cell dysfunction (*PDCD1, HAVCR2, CTLA4, TIGIT*, and *LAG3*; **Fig 2a-b**)^9^. Finally, CD8^+^ T cells also included an intermediate ‘exhausted/activated’ cluster, which contained cells that expressed genes associated with both T cell activation and exhaustion. Notably, comparisons of cluster frequencies pre- and post-therapy revealed an increased frequency of Tfh cells and activated, exhausted, and exhausted/activated CD8^+^ T cells after treatment, supporting reports that PD-1 blockade primarily impacts the expansion of CD8^+^ T cell phenotypes in the TME (**Fig. 2b**)^32, 33^.

**Figure 2.**
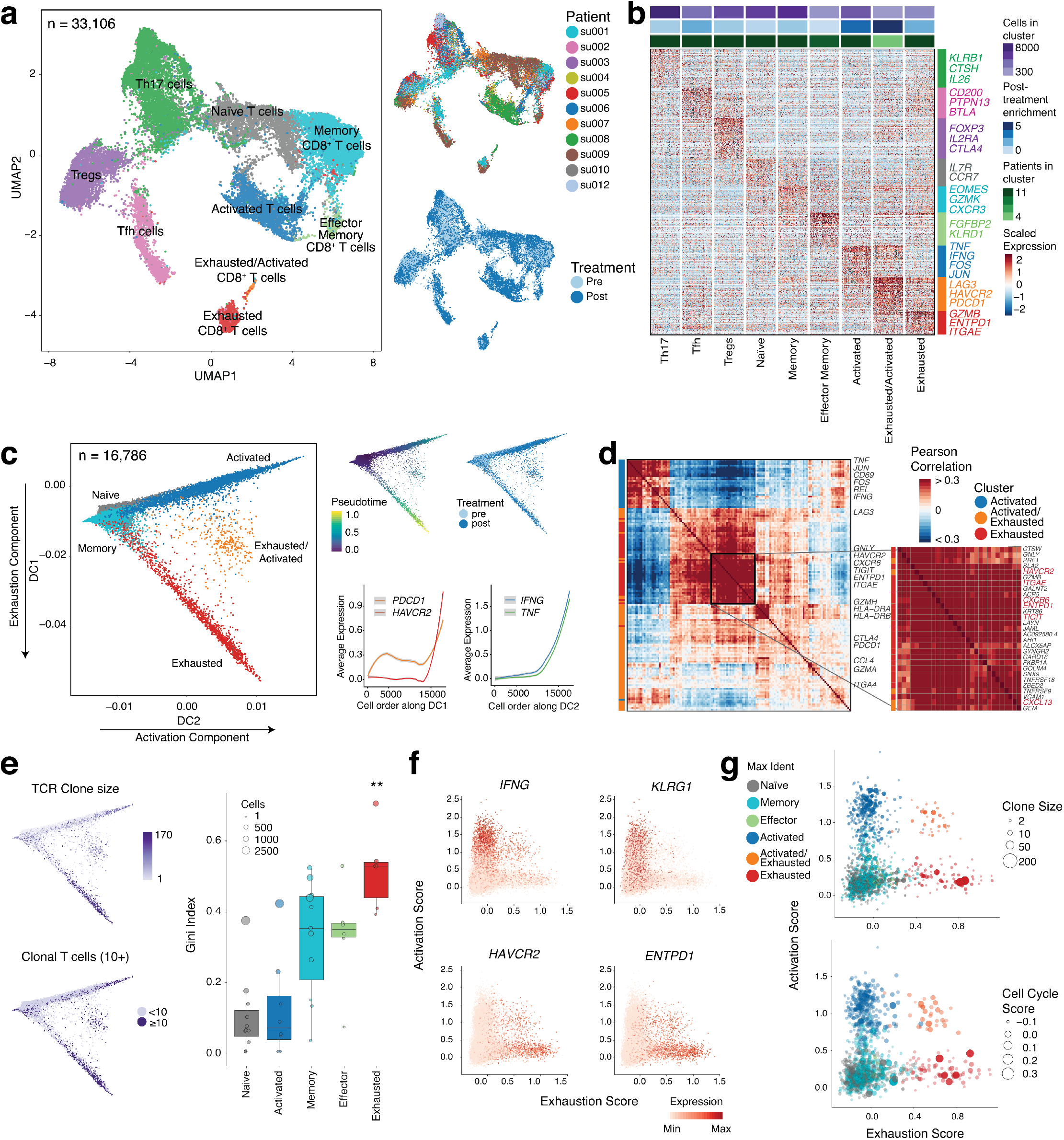
Exhausted CD8^+^ T cells are clonally expanded and express markers of tumor-specificity. **(a)** UMAP of tumor-infiltrating T cells present in BCC samples pre- and post-PD-1 blockade. Clusters denoted by color labeled with inferred cell types (left). UMAP of T cells are also colored by patient (top right) and anti-PD-1 treatment status (bottom right). **(b)** Heatmap of differentially expressed genes (rows) between cells belonging to different T cells subsets (columns). Specific genes associated with different T cell clusters are highlighted. Top bars indicate the number of cells in each cluster, enrichment of clusters pre- and post-therapy, and number of patients in each cluster. **(c)** Diffusion map of naïve, memory, activated and exhausted CD8^+^ T cells using the first two diffusion components (left). Cells colored based on cluster identities from Fig. 2a. Cells are also colored by diffusion pseudotime and treatment status (top right). Expression of selected core activation and exhaustion genes is shown along diffusion components 1 and 2 (bottom right). **(d)** Co-expression analysis of differentially expressed genes between activated, exhausted and activated/exhausted CD8^+^ T cells. Inset indicates a core exhaustion module identified by hierarchical clustering, with canonical exhaustion genes highlighted. **(e)** Diffusion map of CD8^+^ T cell subsets colored by clone size (left) and boxplot of Gini indices for each CD8^+^ T cell cluster (right) calculated for each patient, showing significant clonal expansion within exhausted CD8^+^ T cells (paired t-test, twotailed, \*\**p* < 0.01). **(f)** Activation score (based on expression of top 50 genes most correlated with *IFNG* expression) versus exhaustion score (based on expression of top 50 genes most correlated with *HAVCR2* expression) for all CD8^+^ T cells, colored by expression levels of individual genes. **(g)** Activation score versus exhaustion score for TCR clones based on average activation and exhaustion scores of individual cells belonging to that clone, colored by the most frequent assigned phenotype for cells belonging to that clone, and size based on clone size (top right) or cell cycle score (bottom right).

We used diffusion maps to visualize the relationship between CD8^+^ T cell clusters on a pseudotime trajectory which placed each single cell along a two-component trajectory (**Methods, Fig. 2c**)^34^. The first diffusion component separated activated and exhausted cells, and the second diffusion component separated naïve and memory cells from activated and exhausted cells (**Fig. 2c, Supplementary Fig. 5a**). Accordingly, the first diffusion component was highly correlated with terminal differentiation and T cell exhaustion genes, and the average expression of dysfunction genes, including *PDCD1* and *HAVCR2*, increased along this component (**Fig. 2c, Supplementary Fig. 5b**). The second diffusion component was highly correlated with the expression of T cell activation genes, and the average expression of cytotoxic genes, including *IFNG* and *TNF*, increased along this component (**Fig. 2c, Supplementary Fig. 5b**). To define a core gene signature of T cell exhaustion in the setting of immunotherapy, we analyzed the coexpression of differentially expressed genes between activated and exhausted clusters (**Fig. 2d**). This analysis identified an exhaustion-specific gene expression module containing known exhaustion markers (*HAVCR3, TIGIT*), tissue resident memory T cell (Trm) markers (*ITGAE, CXCR6*)^35, 36^, and the extracellular ATPase CD39 (*ENTPD1*), which has been shown to identify CD8^+^ TILs that recognize tumor antigens^13, 37–39^. The high levels of *PDCD1* (PD-1)^40^, presence of *ITGAE* (CD103) and lack of KLRG1^13, 37–39^ in exhausted clusters also supported an enrichment of tumor-specific T cells in this population. Altogether, these results suggest that exhausted CD8^+^ TILs increase after PD-1 blockade and express gene signatures of chronic activation, T cell dysfunction, and tumor reactivity.

Since tumor antigen-specific CD8^+^ T cell clones will expand during a productive immune response (pre- or post-therapy), we analyzed scTCR-seq profiles to identify clonally-expanded cells as a surrogate marker for tumor-specificity. We grouped cells by clonotype based on both TCRα and TCRβ chain sequences and first examined the distribution of clone sizes found in each cluster (**Fig. 2e**). This analysis showed that the exhausted T cell cluster was enriched for large clone sizes compared to other CD8^+^ clusters. We also calculated the Gini index, a statistical measure of distribution inequality, for each scRNA-seq cell cluster and found that exhausted T cells had significantly higher clonality compared to all other CD8^+^ T cell clusters (mean Gini index = 0.46 for exhausted T cells, mean Gini index = 0.19 for other CD8^+^ clusters; p = 0.0022; **Fig. 2e**). We further examined the relationship between T cell states and clonal expansion by scoring all CD8^+^ T cells for activation and exhaustion signatures, calculated from the top 50 genes correlated with *IFNG* and *HAVCR2* expression, respectively (**Methods, Fig. 2f**). We found that enrichment of the two signatures in individual cells was largely mutually exclusive, and only a small population of cells showed high enrichment of both exhaustion and activation signatures. As in the prior analysis, we found that T cells with a high exhaustion signature enrichment also exhibited patterns of gene expression associated with tumor reactivity, including the expression of CD39 (*ENTPD1*) and CD103 (*ITGAE*), and the absence of *KLRG1* expression^13, 37–39^. To examine the phenotypes of individual expanded clones in this context, we grouped individual cells by clonotype and assigned exhaustion and activation scores to each clone based on the average score of all cells belonging to that clone (**Fig. 2g**). Indeed, the most expanded TIL clones showed a high exhaustion gene signature. We also examined the proliferative capacity of distinct clones by scoring individual cells for expression of cell cycle associated genes (**Methods**) and calculating the average cell cycle score for all cells belonging to each clone. Exhausted cells also exhibited a high proliferation signature, in line with similar analysis of dysfunctional CD8^+^ T cells in melanoma^38^. Of note, clones with the highest exhaustion signature enrichment were not considerably proliferative or clonally expanded, similar to previous reports suggesting that later stages of T cell dysfunction may have limited proliferative potential^38^. Analyses of clonal expansion in CD4^+^ T cell clusters demonstrated a similar increase in clonality in Tfh cells following treatment, which in one patient was accompanied by an increase in B cells expressing germinal center markers, suggesting the presence of a tertiary lymphoid organ (**Supplementary Fig. 6a-d**).

To understand the lineage relationships between T cell phenotypes, we performed a paired analysis of T cell clonotypes and phenotypes. Globally, we found that T cells of the same clonotype were significantly more likely to share a common phenotype and have more highly correlated gene expression signatures than randomly grouped cells (*p* < 2.2 × 10^−16^, unpaired t-test), in line with prior studies (**Fig. 3a-c, Supplementary Fig. 7a-c**)^41–43^. Notably, this property was observed in each individual patient, in CD4^+^ and CD8^+^ T cells, and both pre- and post-therapy (**Fig. 3b, Supplementary Fig. 7a-c**). We also used GLIPH (grouping of lymphocyte interactions by paratope hotspots) to identify ‘TCR specificity groups,’ clusters of distinct TCR sequences that are likely to recognize common antigens via shared motifs in the CDR3 sequence^44^. Similar to the analysis of cells within a given clonotype (that is, sharing identical TCR sequences), T cells expressing distinct TCRs within a specificity group were also more likely to share a common phenotype (pre-treatment: *p* = 0.00035, post-treatment: *p* = 0.0088, unpaired t-test) and have more highly correlated gene expression signatures compared to randomly grouped TCRs (pre-treatment: *p* < 2.2 × 10^−16^, post-treatment: *p* = 3.7 × 10^−16^, unpaired t-test; **Fig. 3a-c, Supplementary Fig. 7d**). However, cells with distinct TCRβ sequences in the same specificity group were significantly more dissimilar than cells which shared the same TCR sequences (*p* < 2.2 × 10^−16^, unpaired t-test; **Fig. 3c**). These results suggest that clonally-expanded TILs are highly correlated in cellular phenotype (perhaps due to the epigenetic stability of parent T cells^11^), and that PD-1 blockade does not lead to instability of phenotypes or the emergence of distinct TIL phenotypes within clones. Moreover, the comparison of specificity groups suggests that antigen context may also have a role in establishing T cell fate choice.

**Figure 3.**
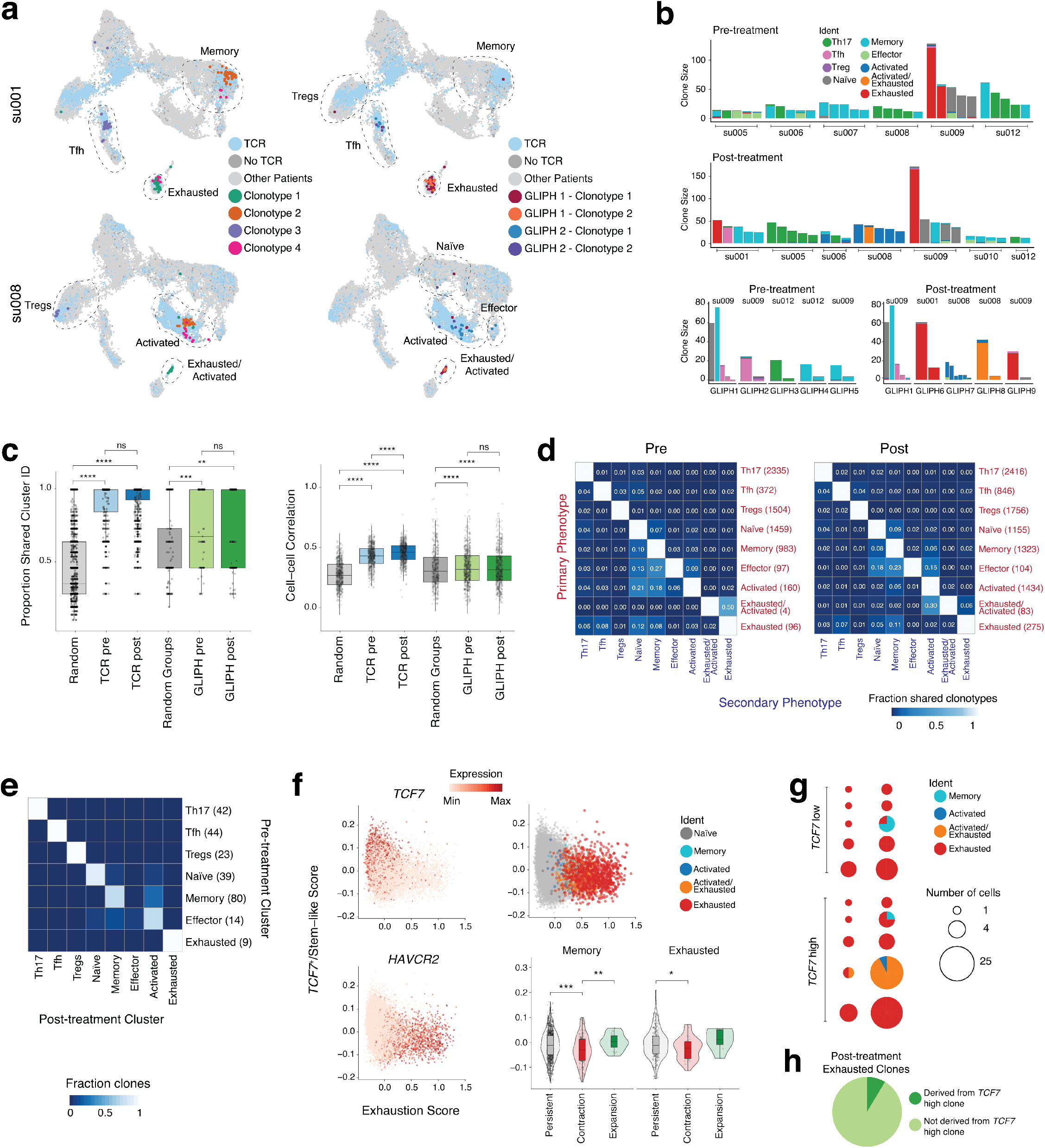
Clonal dynamics and phenotype transitions of tumor-infiltrating T cells. **(a)** UMAP of tumor-infiltrating T cells colored by selected TCR clones (left). UMAP of T cells colored by TCRβ clones belonging to the same TCR specificity group (right). **(b)** Phenotypes of single cells belonging to the same TCR clone or TCR specificity group. Shown are the top five most abundant clones (top and middle) larger than 10 cells for each patient. Each bar is colored by individual phenotypes of single cells within the clone. The bottom plots show phenotypes of distinct TCR clones within a TCR specificity group. Both analyses show substantial phenotypic similarity among single cells belonging to a clone or group. **(c)** Distribution of the proportion of cells within each clone, or TCRβ clones within each TCR specificity group (>=3 cells), that share a common cluster identity, separated by treatment timepoint, compared to randomly selected and size matched groups of T cells from the same sample (left, unpaired t-test, two-tailed, \*\*\*\**p*<0.0001, \*\*\**p*<0.001, \*\**p*<0.01). Distribution of cell-cell correlations between cells that belong to the same TCR clone or cells within the same TCR specificity group but different clonotypes, separated by treatment timepoint and compared to randomly selected and size matched groups of T cells from the same sample (bottom, unpaired t-test, two-tailed, \*\*\*\**p*<0.0001). Cell-cell correlations were calculated using log-transformed expression of differentially expressed genes. **(d)** Heatmap showing the fraction of clonotypes belonging to a primary phenotype cluster (rows) that are shared with other secondary phenotype clusters (columns). **(e)** Heatmap of all observed phenotype transitions for matched clones during PD-1 blockade for clones with at least 3 cells for each timepoint. **(f)** TCF7^+^/stem-like score (from Im *et al*^45^) versus exhaustion score for all CD8^+^ T cells, colored by expression of selected genes (left). TCF7^+^/stem-like score versus exhaustion score for exhausted cells and cells of other phenotypes belonging to primarily exhausted clones, colored by phenotype (top right). Violin plot of TCF7^+^/stem-like score for memory and exhausted cells separated by change in clone abundance following treatment (bottom right). Clones were defined as expanded or contracted if they significantly changed in abundance by a Fisher exact test, and persistent if they did not significantly change in abundance and at least one cell detected at each timepoint. **(g)** Pie charts showing clone size and distribution of phenotypes across timepoints for matched clones pre- and posttherapy. Clones were only considered if they demonstrated an exhausted phenotype pretherapy and increased in abundance post-therapy. The plot shows clones separated by the presence of a high *TCF7*^+^ signature prior to treatment. **(h)** Pie chart of all clones with an exhausted phenotype post-treatment. Shading indicates whether a *TCF7*^+^ cell of the same clonotype was present pre-therapy.

Since some clones contained cells with divergent phenotypes, we asked whether these cells could indicate lineage transitions between T cell phenotypes. We aggregated all clonotypes belonging to a given cluster (‘primary phenotype’) and measured the fraction of clonotypes that belonged to another cell cluster (‘secondary phenotype’) (**Fig. 3d**). Broadly, we noted significant overlaps between either activated, memory, effector memory, or exhausted CD8^+^ T cell phenotypes, demonstrating that CD8^+^ T cells clusters represent a continuum of activation and memory states that can be accessed by cells within a clone. For example, primarily activated T cell clones showed the most overlap with naïve and memory T cells, suggesting a transition between memory and activated states. Similarly, primarily exhausted or exhausted/activated cells showed overlaps with each other, or with memory or activated cells, respectively. In line with recent reports, we detected minimal clonotype sharing between exhausted and effector cells, suggesting a strict bifurcation in the fate decision towards either phenotype^38^. In contrast to CD8^+^ T cells, CD4^+^ T cell clones were largely restricted to single phenotypes, suggesting minimal plasticity of cell states. For example, Treg clonotypes were almost exclusively restricted to the Treg cell fate (**Fig. 3d**). We also examined divergent phenotypes within specificity groups by comparing the distribution of phenotypes for individual clones within specificity groups (**Supplementary Fig. 7e**). We found that this distribution was non-random, suggesting that certain groups of T cell phenotypes may be similarly driven by specific TCR signal strength thresholds. For example, specificity groups containing exhausted clones are most often shared with clones of memory and naïve phenotypes, but never with clones of effector phenotypes. Unlike phenotype overlap within TCR clones, we noted more significant overlaps between CD8^+^ and CD4^+^ phenotypes within specificity groups, such as the co-occurrence of CD8^+^ exhausted and CD4^+^ Treg and Tfh clones within the same specificity group. These results suggest that CD4^+^ and CD8^+^ TILs responding to the same antigen may arise from distinct clonotypes (**Supplementary Fig. 7e**).

To track clonal cell fates after PD-1 blockade, we matched clonotypes between treatment timepoints, based on shared TCRβ sequences, and detected on average 121 overlapping clonotypes present at any frequency between paired patient samples (range 6-531, median 65), with the exception of one pair with limited cell numbers and no clonotype overlap (su003) (**Supplementary Fig. 8a**). We detected on average <1 overlapping clone between different patients, confirming the accuracy of tumor sample identities. To globally assess treatment-induced clone transitions across all patients, we compared the primary phenotypes at each timepoint for matched clones with at least three cells per timepoint (**Fig. 3e, Supplementary Fig 8b**). We observed stability among CD4^+^ T cell clusters and frequent transitions among CD8^+^ T cell clusters, similar to the pattern observed in individual timepoints. While we observed frequent transitions between memory and effector states to an activated state post-treatment, clones in an exhausted state prior to treatment did not transition to non-exhausted phenotypes post-treatment, suggesting that exhausted T cells may be limited in their capacity for phenotype transition after PD-1 blockade.

Prior studies identified a stem-like T cell population expressing the transcription factor *TCF7* that provides the proliferative T cell burst in response to PD-1 blockade^33, 45–48^. We asked whether similar cells could be observed in our dataset, and whether these cells persisted in post-therapy samples. We scored individual CD8^+^ cells for the expression of exhaustion signature genes or *TCF7*^+^ stem-like signature genes, as defined previously (**Fig. 3f**)^45^. This analysis revealed three primary categories of cells: 1) cells with a high *TCF7*^+^ signature and low exhaustion signature, consisting primarily of naïve, activated and memory cells, 2) cells with a high exhaustion signature and low *TCF7*^+^ signature, consisting primarily of exhausted and activated/exhausted cells, and 3) cells with low expression of both signatures. We further examined all cells belonging to exhausted clones and observed a small population of exhausted cells (28% of exhausted cells, 1.5% of all CD8^+^ T cells) with expression of both *TCF7*^+^ and exhaustion signatures (**Fig. 3f**). Since these cells have been shown to proliferate after PD-1 blockade, we asked whether clones that persisted or expanded post-therapy in each patient displayed a higher *TCF7*^+^ signature pre-therapy compared to clones that contracted. Indeed, we found that for both memory and exhausted phenotypes, persistent clones had a significantly higher *TCF7*^+^ signature pre-treatment compared to clones that contracted (**Fig. 3f**). However, we only identified two exhausted clones that significantly expanded following treatment, limiting our ability to assess their *TCF7*^+^ signature expression pre-therapy. To look into this further, we identified 10 exhausted clones pre-treatment that increased in frequency posttherapy but were excluded from the previous analysis due to low clonal size and limited expansion (**Fig. 3g**). Of these clones, the majority remained exhausted after therapy, but we did observe that those with a high *TCF7*^+^ signature pre-treatment expanded more substantially than those with a low *TCF7*^+^ signature, confirming the prior proliferative behavior of these cells in the setting of PD-1 blockade. Nevertheless, this phenomenon only appeared to explain a minority of T cell expansion post-therapy, since only a small fraction (10%) of post-therapy clones were derived from exhausted clones containing *TCF7*^+^ cells pre-therapy, suggesting that post-treatment exhausted clones may be derived from additional sources (**Fig. 3h**).

Since few pre-existing exhausted T cells showed expansion post-therapy, we asked how clone abundance changed globally in response to treatment by comparing the pre-treatment frequency of each clone to its frequency post-treatment based on scTCR-seq (**Fig. 4a**). We noted that while the majority of clones did not significantly change in abundance (93%), we could detect many significantly expanded clones post-therapy that were not detected prior to treatment (68% of significantly expanded clones). Integration of scRNA-seq data revealed strikingly different patterns of persistence for each cell phenotype. Namely, compared to other CD8^+^ T cell subsets, a relatively small fraction of post-therapy exhausted T cell clones could be observed anywhere in the pre-therapy tumor; on average only 40% of naïve, activated, memory, or effector memory clones were derived from novel clonotypes, while 84% of exhausted clones were derived from novel clonotypes (p = 0.027, unpaired t-test, **Fig. 4a-b**). We further visualized the contribution of pre-existing clones to overall clonality of each population using a Lorenz curve, which shows the cumulative proportion of all TCRβ clones ranked by size compared to the cumulative proportion of the T cell subset covered by those clones (**Fig. 4c**). Among memory T cells post-therapy, only 20% of expanded clones greater than or equal to 5 cells in size were derived from novel clonotypes. In contrast, 55% of the top clones in the exhausted cluster post-therapy were novel. We next asked how the expansion of novel clones, a phenomenon we termed ‘clonal replacement,’ contributed to the overall frequency of exhausted T cells in each patient (**Fig. 4d, Supplementary Fig. 9a**). We found that 7/11 patients had an increased frequency of exhausted CD8^+^ cells following treatment, and in 6/7 patients, the majority of post-treatment exhausted cells were derived from novel clonotypes (**Fig. 4d**). We also compared the proportion of exhausted cells post-treatment in responder and non-responder patients and noted some differences between groups (**Fig. 4d**). However, meaningful associations of clonal replacement with clinical outcomes will require further work in much larger and prospective studies across diverse patient populations. These findings suggest that PD-1 blockade does not act by re-invigorating exhausted tumor-specific T cells that are clonally-expanded prior to therapy, but rather by expanding a distinct TCR repertoire. Interestingly, novel posttreatment exhausted clones were enriched for novel TCR specificity groups, suggesting that novel clones may also represent new antigen specificities (**Supplementary Fig. 9b**)

**Figure 4.**
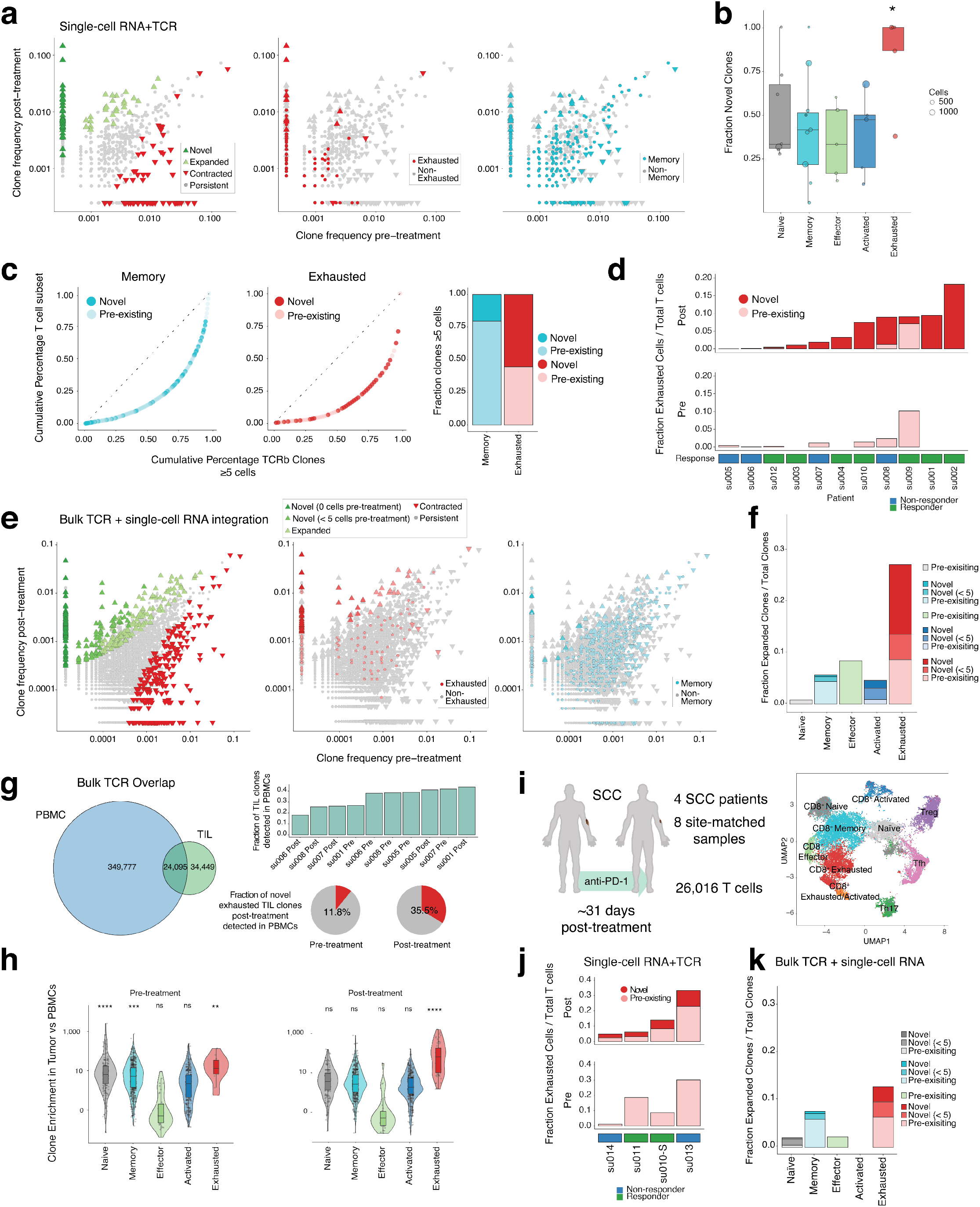
Clonal replacement of CD8^+^CD39^+^ T cells following PD-1 blockade. **(a)** Scatterplots comparing TCRβ clone frequencies pre- and post-treatment measured by single-cell RNA_+_TCR-seq for all BCC patients (n =11 patients). Clones that were significantly expanded or contracted post-treatment based on a Fisher exact test are highlighted on the left. Clones where the majority of cells exhibit an exhausted CD8^+^ phenotype (middle, red) or a memory CD8^+^ phenotype (right, blue) are also highlighted. (b) Boxplot of the fraction of novel clones detected by scRNA_+_TCR-seq within each cluster following treatment (unpaired t-test, two-tailed, \**p*<0.05). Clones with only one cell detected in total and cells from su003 with no clonotype overlap between timepoints were excluded. **(c)** Lorenz curve of TCRβ clone frequencies based on scRNA_+_TCR-seq for exhausted CD8^+^ T cell clones (left) and memory CD8^+^ T cell clones (middle) greater than or equal to 5 cells, colored by presence of each clone prior to treatment. Proportion of novel clones in each phenotype quantified on the right. **(d)** Fraction of exhausted cells out of total T cells detected by single cell RNA_+_TCR-seq for each patient, separated by treatment status. Cells belonging to novel clones detected post-treatment are highlighted. **(e)** Scatterplots comparing TCRβ clone frequencies pre- and post-treatment measured by bulk TCR-seq (n = 8 patients). Clones that were significantly expanded or contracted post-treatment based on a binomial test are highlighted on the left, with expanded clones further separated based on their detection pre-treatment. Clones where the majority of cells share an exhausted CD8^+^ phenotype based on scRNA-seq (middle, red) or a memory CD8^+^ phenotype (right, blue) are also highlighted. **(f)** Bar plot of fraction of clones with significant expansion post-treatment based on bulk TCR-seq, separated by phenotype and colored by replacement status. **(g)** Overlap between TCRβ clones in peripheral blood and tumor infiltrating T cells detected by bulk TCR-seq (n = 5 patients, left). Fraction of TIL clones detected in peripheral blood, separated by sample (top right). Fraction of novel exhausted TIL clones detected in PBMCs, separated by treatment status (bottom right). **(h)** Violin plot of clone enrichment (tumor frequency / PBMC frequency) detected by bulk TCR-seq, separated by phenotype and treatment status (n = 5 patients, unpaired t-test, one-tailed, \*\*\*\**p*<0.0001, \*\*\**p*<0.001, \*\**p*<0.01, \**p*<0.05). **(i)** Characteristics of squamous cell carcinoma (SCC) samples treated with anti-PD-1 (left) and UMAP of tumor-infiltrating T cells present in SCC samples pre- and post-PD-1 blockade (right). Clusters denoted by color are labeled with inferred cell types. **(j)** Fraction of exhausted cells out of total T cells detected by single-cell RNA_+_TCR-seq for each patient, separated by treatment status. Novel clones detected post-treatment are highlighted (bottom left). Sample su010-S derived from an SCC lesion from patient su010 who presented with both BCC and SCC lesions. **(k)** Bar plot of fraction of clones with significant expansion based on bulk TCR sequencing post-treatment, separated by phenotype and colored by replacement status (bottom right).

To increase sampling depth and sensitivity for detecting rare clonotypes, we performed bulk TCR-seq on the remaining biopsy material from 8/11 patients (**Methods**). We obtained on average 19,114 TCR templates from tumor samples (range 554-71,031), representing an average of 5,527 unique clonotypes (range 237-14,197) per sample, over a 20-fold increase in TCR sampling depth compared to the scRNA-seq experiments. We again compared pre- and post-therapy TCR sequences from each patient and identified clones that expanded or contracted post-therapy, excluding clones with less than five templates detected (**Fig. 4e**). Similar to the analysis done with scTCR-seq data, we observed a significant number of novel expanded clones that were either not expanded prior to treatment (<5 cells) or entirely absent pre-treatment, and these clones were enriched in cells annotated with an exhausted phenotype based on integration of scRNA-seq data. Indeed, compared to all other CD8^+^ phenotypes, exhausted cells had a higher proportion of significantly expanded clones following treatment, and the majority of expanded clones were derived from novel clonotypes (**Fig. 4f**). Again, clonal replacement of exhausted clones was observed in every patient that demonstrated expanded exhausted cells post-therapy (**Supplementary Fig. 9c**). Conversely, in non-exhausted CD8^+^ T cell populations with significantly expanded clones following treatment, across all patients, few were derived from novel clones (**Supplementary Fig. 9c**). To address whether these effects could simply be attributed to sampling bias rather than PD-1 blockade effects, we sampled one patient twice before therapy (site-matched but at different timepoints) and twice after therapy (site-matched but at different timepoints) at approximately two-month intervals (**Supplementary Fig. 9d**). Importantly, we only observed clonal replacement of exhausted clones when comparing pre-to post-treatment samples, but not when comparing two timepoints within pre- or post-treatment groups, suggesting that TCR dynamics of exhausted cells were mainly influenced by PD-1 blockade and not the timing or location of tumor biopsies. Altogether, these results confirm that after PD-1 blockade, a significant proportion of clonally-expanded CD8^+^CD39^+^ T cells arise from novel clones, which are not detected in the tumor prior to treatment.

We next asked whether novel clonally-expanded T cells in the tumor could also be detected in the peripheral blood. We performed bulk TCR-seq on 12 peripheral blood samples taken at the time of tumor biopsy either pre- or post-therapy, with 10 peripheral blood samples from 5 patients also matched with bulk TCR-seq from TILs. Overall, 41% of TIL TCR clonotypes could also be detected in blood (range: 18-44% per sample), but only represented 6% (range 0.2-19%) of the total number of blood clonotypes (**Fig. 4g, Supplementary Fig. 10a**). Importantly, blood clonotypes represented clones from all T cell phenotypes identified in the tumor, and in all patients who demonstrated a significant expansion of exhausted clones in the tumor (> 5 exhausted clones detected by scRNA-seq), these cells could also be detected in the blood (**Supplementary Fig. 10a,b**). Moreover, novel clonally-expanded clonotypes in the tumor post-therapy could be detected in the peripheral blood both post- and pre-therapy, suggesting that peripheral sources of T cells might be recruited to drive the response to PD-1 blockade (**Supplementary Fig. 10a**). Overall, 35.5% of novel exhausted clonotypes detected posttreatment in the tumor could also be detected in the peripheral blood post-therapy, suggesting that clones responding to PD-1 blockade may circulate in both sites. Surprisingly, 11.8% of novel exhausted clonotypes detected in the tumor post-treatment could even be detected in peripheral blood pre-treatment, demonstrating that clones which are undetectable by deep TCR sequencing in the tumor prior to treatment could be detected in the peripheral blood (**Fig. 4g**). We also compared clonotype enrichment in the tumor to the peripheral blood for different phenotypes by comparing the frequency of each clonotype in each location. Many CD8^+^ phenotypes showed a significant enrichment in the tumor compared to the peripheral blood, but we noted a significant increase in the enrichment of exhausted clones compared to other phenotypes following treatment, suggesting that these clones may preferentially expand and be retained in the tumor, supporting their tumor-specificity (*p* < 2×10^−16^, unpaired t-test, **Fig. 4h**). We observed a similar trend for an enrichment of exhausted TCR specificity groups compared to other phenotypes in the TME following treatment, suggesting that antigen-specificity drives the expansion of these cells in the tumor (**Supplemental Fig. 10c**). These results suggest that it may be feasible, though challenging, to monitor the clonal tumor-specific T cell response to checkpoint blockade in the blood^49, 50^.

Finally, we asked whether clonal replacement of exhausted cells could be observed in a different cancer type. We generated scRNA_+_TCR-seq profiles of 26,016 tumor-infiltrating T cells from serial tumor biopsies in 4 patients with SCC treated with anti-PD-1 (**Fig. 4i, Supplementary Fig. 11a**). SCC is a natural choice for comparison to BCC, since this cancer type also exhibits high levels of UV-induced tumor mutation burden, is accessible for site-matched pre- and post-therapy samples, and has a similar clinical response rate to PD-1 blockade^51^. Due to clinical constraints, SCC samples were obtained an average of 31 days post-treatment, enabling an analysis of TIL clonotype/phenotype dynamics relatively early after treatment. We first confirmed a number of our prior findings in the context of SCC: 1) TIL phenotypes in SCC were highly correlated TIL phenotypes in BCC (**Supplementary Fig. 11b**), 2) exhausted CD8^+^ T cells were clonally expanded (**Supplementary Fig. 11c**) and marked by surface proteins associated with tumor-specificity, including CD39^13, 37–39^ (**Supplementary Fig. 11d**), 3) TILs sharing a TCR clonotype or TCR specificity group were highly correlated in cellular phenotype, and PD-1 blockade did not lead to phenotype instability or the emergence of new phenotypes (**Supplementary Fig. 11e-f**), 4) T cell clone phenotypes were largely stable between treatment timepoints (**Supplementary Fig. 11g**), and 5) clones with a high *TCF7*^+^ signature were more likely to persist after therapy (**Supplementary Fig. 11h,i**). Next, we compared clonal replacement in pre- and post-therapy SCC samples and observed that a significant proportion of exhausted cells detected post-treatment were derived from novel clonotypes (**Fig. 4j**). Integration of scRNA-seq data with bulk TCR-seq confirmed that novel clones comprised a significantly larger portion of expanded clones compared to other phenotypes, with overall only 29% of expanded naïve, activated, memory, or effector memory clones derived from novel clonotypes, while 50% of expanded exhausted clones were derived from novel clonotypes (**Supplementary Fig. 11j, Fig. 4k**). Notably, we observed a similar degree of expansion of pre-existing clones at early timepoints in SCC as we did at later timepoints in BCC, suggesting that pre-existing clones have a limited capacity for clonal expansion and that the T cell response to PD-1 blockade is primarily derived from novel clones.

## Discussion

Here, we performed single-cell multi-omic profiling of clinical tumor biopsies to measure the clonal evolution of the T cell response to PD-1 blockade. Integration of TCR clonotype and scRNA-seq phenotype in 51,493 single TILs revealed several principles of the tumor-specific T cell response. First, although the TME contained a rich diversity of T cell phenotypes, including several CD4^+^ T helper subtypes and a spectrum of CD8^+^ activation and dysfunction phenotypes, clonally-expanded cells post-therapy were highly-enriched in CD8^+^ T cells that were phenotypically exhausted. These cells co-expressed several markers of T cell dysfunction, as well as markers previously shown to enrich for tumor-specific T cells, including CD39 and CD103^13, 37–39^. These results suggest that tumor recognition, clonal expansion, and T cell dysfunction are intimately linked processes, and that the TME contains a large number of ‘bystander’ T cells, as has been previously suggested^13, 52^. Second, tracking individual T cell clones before and after therapy in site-matched specimens provided insights into the origin of clonally-expanded T cells. Namely, while a significant fraction of memory CD8^+^ T cell clones persisted in tumors post-therapy (as did a similar fraction of all other non-exhausted clones), the clonal repertoire of exhausted CD8^+^ T cells was largely replaced by novel clones after therapy. This phenomenon was observed in both BCC and SCC patients and was confirmed by deep TCR sequencing of tumor biopsies. Taken together, these results support the concept that the chronic activation and exhaustion of pre-existing tumor-infiltrating T cells limits their re-invigoration following checkpoint blockade^11^, and that the T cell response relies on the expansion of a distinct repertoire of tumor-specific T cell clones.

Clonal replacement of the tumor-specific T cell response is consistent with several prior lines of investigation. First, studies in mouse models of chronic viral infection have demonstrated that T cell exhaustion is associated with a broad remodeling of the epigenetic landscape, which underlies the inability to stably reverse this phenotype through PD-1 blockade^11, 12, 53^. Second, analysis of T cells that proliferate after PD-1 blockade during viral infection identified a population of responding CD8^+^ cells which expressed markers associated with Tfh cells, such as CXCR5^45^. Notably, CXCR5^+^CD8^+^ cells were only present in lymphoid organs, and not in tissues, suggesting that the response to PD-1 blockade is initiated outside of tissues and requires migration to the site of infection or cancer. Accordingly, studies in murine cancer models demonstrated that checkpoint inhibitors initiated a systemic immune response in many organs, and that chemically blocking T cell migration abrogated the anti-tumor T cell response^54, 55^. It is important to note that our study did not definitively identify the organ source of novel expanded T cell clones post-therapy, and at least two possibilities exist: 1) novel clones could originate from tumor-extrinsic sources, such as lymphoid organs, or 2) novel clones could originate from rare unexpanded clones present within the TME or tumor periphery. Although the bulk TCR sequencing results support the first possibility, this study is limited by the sampling (time and material) constraints of human tumor biopsies, and further work will be required to dissect the timing and extent of pre-existing TIL expansion after PD-1 blockade in human tumors. Importantly, both possibilities are compatible with the potential derivation of these cells from a TCF7^+^ precursor T cell population, which has been shown to proliferate after PD-1 blockade (both in the tumor and in peripheral sites) and correlates with clinical outcomes^33, 45–48^.

Regardless of the site of origin, our results demonstrate that PD-1 blockade unleashes a novel tumor-specific TCR repertoire that was not previously expanded in the tumor, which has significant implications for patient prognosis and treatment. Namely, efforts to predict the response to immunotherapy from blood may be more informative than from the pretreatment tumor, and efforts to expand TILs for therapy may need to identify relevant clones rather than bystander cells in the tumor. These findings also raise several questions. First, how do the clonal origins of T cell response differ in CTLA4 blockade, and how does this impact PD-1 blockade? It is possible that these agents recruit distinct phenotypic populations of novel clones to the tumor, which have synergistic effects. Second, the presence of novel clones suggests an alternative hypothesis for the improved activity of checkpoint blockade agents in immune-infiltrated (‘hot’) vs immune-desert (‘cold’) tumors. That is, perhaps the reason that ‘hot’ tumors respond to therapy is due to an intrinsic ability of the tumor to constantly attract new T cells^56^, rather than the presence of pre-existing T cells in the tumor. Therefore, therapeutic agents that improve recruitment or infiltration of T cells into the tumor might be sufficient to rescue therapeutic responses in cold tumors. Finally, the expansion of novel TCR clones and TCR specificity groups following PD-1 blockade, coupled with the presence of neoepitope loss, suggests that pre-existing and novel T cell clones may recognize different tumor antigens and initiate distinct waves of immunoediting^57^. The identity of antigens recognized by each wave and the biophysical properties of their TCR:peptide-MHC interactions require further investigation, perhaps using high-throughput tumor specificity assays^52, 58^. In summary, this study reveals new insights into the clonal T cell response to checkpoint blockade in human cancer, which has important implications for the future design of checkpoint blockade immunotherapies.

## Acknowledgments

We thank members of the Chang laboratory for helpful discussions. We thank X. Ji, D. Wagh, and J. Coller at the Stanford Functional Genomics Facility, J. Sung and S. Fitch at the Stanford Clinical and Translational Research Unit Biobank, and D.D. Hiraki at the Stanford Blood Center Histocompatibility, Immunogenetics, and Disease Profiling Laboratory. We thank A.D. Colevas, S. Reddy, L. Van Der Bokke, R. Patel for clinical collaboration. We thank A. Pague and A. Valencia for assistance with clinical specimens. We thank Khriszha Quema-Yee for assistance with illustrations. This work was supported by the Parker Institute for Cancer Immunotherapy (A.T.S., D.K.W., R.K., S.L.B., M.M.D., and H.Y.C.), the Michelson Foundation (A.T.S.), and the National Institutes of Health (NIH) P50HG007735, R35CA209919 (H.Y.C.), K08CA23188-01 (A.T.S.), K23CA211793 (K.Y.S.) and 5U19AI057229 (M.M.D.). K.E.Y. was supported by the National Science Foundation Graduate Research Fellowship Program (NSF DGE-1656518) and a Stanford Graduate Fellowship. A.T.S. was supported by a Bridge Scholar Award from the Parker Institute for Cancer Immunotherapy, a Career Award for Medical Scientists from the Burroughs Wellcome Fund, and the Human Vaccines Project Michelson Prize for Human Immunology and Vaccine Research. Cell sorting for this project was done on instruments in the Stanford Shared FACS Facility. Sequencing was performed by the Stanford Functional Genomics Facility (supported by NIH grant S10OD018220). M.M.D. and H. Y.C. are investigators of the Howard Hughes Medical Institute.

## Author Contributions

K.E.Y., A.T.S., A.L.S.C. and H.Y.C. conceived the project. K.E.Y., A.T.S., Y.Q., R.K.G., R.A.B. and K.Y.S. performed experiments. K.E.Y., A.T.S., D.K.W., R.K., C.W., K.M., J.M.G., R.A.B., and K.Y.S. analyzed data. S.B., C.C., M.M.D., A.L.S.C., and H.Y.C. guided data analysis. K.E.Y., A.T.S., and H.Y.C. wrote the manuscript with input from all authors.

## Methods

### Human subjects

This study was approved by the Stanford University Administrative Panels on Human Subjects in Medical Research, and written informed consent was obtained from all participants. Patients were treated with 200 mg pembrolizumab every 3 weeks or 350 mg cemiplimab every 2 weeks. A subset of patients received ongoing treatment with 150 mg vismodegib daily (Supplementary Table 1). Response was assessed by RECIST version 1.1^14^.

### Sample Collection and Processing

Fresh biopsies were collected from the primary tumor site. A portion of the tumor was stored in RNAlater for whole exome sequencing and bulk TCR sequencing. The remaining tissue was processed for single cell RNA sequencing.

### H&E and Immunohistochemical (IHC) staining

For H&E staining, formalin-fixed, paraffin-embedded tissue cut at 4 microns and stained using the Tissue-Tek Prisma automated slide stainer. Immunostaining was performed on the Ventana Benchmark Ultra platform (CD3) and the Leica Bond platform (CD8, PD-L1). Antibodies used include anti-human CD3 (cat. no. 103A-76, Cell Marque), anti-human CD8 (cat. no. M7103, Dako) and anti-human PD-L1 (cat. no. 13684S, Cell Signaling Technology).

### Exome Sequencing

Exome capture was performed by Accuracy and Content Enhanced (ACE) augmented exome strategy (Personalis) and sequenced on an Illumina HiSeq 2500 with paired-end 100-bp sequencing, with an average of 110-fold coverage (range 82-138).

### HLA Typing

All samples were genotyped for HLA-A, -B, -C, -DPA1, -DPB1, -DQA1, -DQB1, -DRB1 and -DRB3/4/5 loci using the MIA FORA NGS FLEX HLA Typing 11 Kit 96 Tests (Immucor, Inc. Norcross, GA, USA), following manufacturer’s semi-automated protocol and as described previously^59, 60^. Briefly, paired-end sequence reads were generated using an NGS-based HLA genotyping method targeting 11 HLA genes with extensive coverage of the HLA genomic region by long-range polymerase chain reaction (PCR). Coverage for HLA class I loci is >200 bp 5’UTR to 3’UTR ~200-400 bp. In the case of HLA-DQA1 locus is ~200 bp of the 5’UTR to ~200 bp of the 3’UTR and for HLA-DQB1 locus is ~70bp of the 5’UTR to ~100 bp of the 3’UTR. For the remaining class II loci specific key regions of the gene were amplified. For HLA-DPA1 locus, this coverage is from exon 1 to exon 4 and for HLA-DPB1 locus from exon 2 to exon 4. All HLA-DRB1/3/4/5 genes were co-amplified in two separate reactions. The coverage for HLA-DRB1/3/4 loci included ~300-500 bp of the 5’UTR to the first ~270 bp of intron-1 and the 3’ end of intron-1 (~250 bp) to exon-6. For the HLA-DRB5 gene exon 2 to exon 6 were amplified. All libraries were sequenced on an Illumina MiSeq. For assignment of HLA genotypes, NGS paired-end reads were analyzed using the MIA FORA FLEX v3.0 software (Immucor). Final HLA genotyping calls were confirmed by manual review.

### Bulk TCR Sequencing

Deep sequencing of the TCRβ gene was performed using the immunoSEQ platform (Adaptive Biotechnologies) on genomic DNA extracted from tumor biopsies or peripheral blood with input amounts ranging from 616 ng to 6,004 ng. Only data from productive rearrangements was exported from the immunoSEQ Analyzer for further analysis. On average, 26,066 TCR templates were detected from tumor samples (range 554-99,264), representing an average of 6,041 unique clonotypes (range 237-17,181), ~20-fold increase in sampling depth compared to scTCR-seq. For peripheral blood samples, on average 113,528 TCR templates were detected (range 24,679-257,772), representing an average of 36,536 unique clonotypes (range 7,041-71,462).

### Tumor Dissociation

Fresh tumor biopsies were minced and digested in 5 mL digestion media (DMEM/F12 media + 250 μg/mL Liberase TL + 200 U/mL DNAse I) in a C-tube using the gentleMACS Octo system at 37°C for 3 hours at 20 rpm. Following digestion, 50 μL of 500 mM EDTA was added and sample collected by centrifugation at 300xg for 5 minutes. The cell suspension was then passed through a 70 μm filter and pelleted by centrifugation at 300xg at 4°C for 10 minutes. Cells were then resuspended in 1 mL of RPMI media and cryopreserved in FBS supplemented with 10% DMSO until further processing.

### Cell Sorting

Cells were categorized as peri-tumoral T cells (CD45^+^CD3^+^), other peri-tumoral lymphocytes (CD45^+^CD3^-^) and tumor/stromal cells (CD45^-^CD3^-^). For patients su009, su010, su011, su012, su013, and su014, only peri-tumoral T cells were isolated and used for scRNA-seq. For sample su010-S, we additionally sorted CD8^+^CD39^+^ peri-tumoral T cells (CD45^+^ CD3^+^ CD8^+^ CD39^+^). Antibodies used included anti-human CD45 conjugated to V500 (clone HI30, cat. no. 560779, BD Biosciences), anti-human CD3 conjugated to FITC (clone OKT3, cat. no. 11-0037-41, Invitrogen), anti-human CD8 conjugated to Pacific Blue (clone 3B5, cat. no. MHCD0828, Invitrogen), anti-human CD39 conjugated to APC (clone A1, cat. no. 328210, BioLegend), anti-human PD-1 conjugated to APC/Cy7 (clone EH12.2H7, cat. no. 329921, BioLegend) and anti-human HLA-DR conjugated to eVolve 605 (clone LN3, cat. no. 83-9956-41, Affymetrix-Ebioscience). For bulk RNA-seq datasets, CD4^+^ T helper cells were sorted as naive T cells (CD4^+^CD25^-^CD45RA^+^), Treg (CD4^+^CD25^+^IL7R^lo^), Th1 (CD4^+^CD25^-^IL7R^hi^CD45RA^-^CXCR3^+^CCR6^-^), Th2 (CD4^+^CD25^-^ IL7R^hi^CD45RA^-^CXCR3^-^CCR6^-^), Th17 (CD4^+^CD25^-^IL7R^hi^CD45RA^-^CXCR3^-^CCR6^+^), Th1-17 (CD4^+^CD25^-^IL7R^hi^CD45RA^-^CXCR3^+^CCR6^+^), and Tfh subsets (CXCR5^+^ counterparts of each). All antibodies were used at a 1:200 dilution, with the exception of anti-CD45 and anti-HLA-DR antibodies which were used at a 1:100 dilution. Propidium iodine (cat. no. P3566, Invitrogen) was used for live/dead staining at a final concentration of 2.5 μg/mL.

### Single-cell RNA-seq Library Preparation

Single-cell RNA-seq and TCR-seq libraries were prepared using the 10X Single Cell Immune Profiling Solution Kit, according to the manufacturer’s instructions. Briefly, FACS sorted cells were washed once with PBS + 0.04% BSA and resuspended in PBS + 0.04% BSA to a final cell concentration of 100-800 cells/μL as determined by hemacytometer. Cells were captured in droplets at a targeted cell recovery of 500-7000 cells, resulting in estimated multiplet rates of 0.4-5.4%. Following reverse transcription and cell barcoding in droplets, emulsions were broken and cDNA purified using Dynabeads MyOne SILANE followed by PCR amplification (98°C for 45 sec; 13-18 cycles of 98°C for 20 sec, 67°C for 30 sec, 72°C for 1 min; 72°C for 1 min). Amplified cDNA was then used for both 5’ gene expression library construction and TCR enrichment. For gene expression library construction, 2.4-50 ng of amplified cDNA was fragmented and end-repaired, double-sided size selected with SPRIselect beads, PCR amplified with sample indexing primers (98°C for 45 sec; 14-16 cycles of 98°C for 20 sec, 54°C for 30 sec, 72°C for 20 sec; 72°C for 1 min), and double-sided size selected with SPRIselect beads. For TCR library construction, TCR transcripts were enriched from 2μL of amplified cDNA by PCR (primer sets 1 and 2: 98°C for 45 sec; 10 cycles of 98°C for 20 sec, 67°C for 30 sec, 72°C for 1 min; 72°C for 1 min). Following TCR enrichment, 5-50 ng of enriched PCR product was fragmented and end-repaired, size selected with SPRIselect beads, PCR amplified with sample indexing primers (98°C for 45 sec; 9 cycles of 98°C for 20 sec, 54°C for 30 sec, 72°C for 20 sec; 72°C for 1 min), and size selected with SPRIselect beads.

### Sequencing

Single-cell RNA libraries were sequenced on an Illumina NextSeq or HiSeq 4000 to a minimum sequencing depth of 25,000 reads/cell using the read lengths 26bp Read1, 8bp i7 Index, 98bp Read2. Single-cell TCR libraries were sequenced on an Illumina MiSeq or HiSeq 4000 to a minimum sequencing depth of 5,000 reads/cell using the read lengths 150bp Read1, 8bp i7 Index, 150bp Read2.

### Data Processing of exome libraries

Whole exome sequencing was preprocessed using a standard GATK approach^61^. Briefly, both tumor and normal samples were aligned to GRCh37 using bwa-mem^62^ and further processed to remove duplicates and recalibrate base quality scores. All processing was performed in FireCloud^63^.

### Mutation calling and neoepitope prediction

Small somatic variants were identified using Mutect2^64^ and further annotated with the GATK. Somatic copy number variants were identified using the GATK best practices pipeline. HLA typing was performed on the germline whole exome sample using xHLA^65^. Neoepitopes were identified using pVAC-seq^17^, where a peptide-MHC pair was considered a neoepitope if the peptide was found to bind to the MHC allele with less than 500 nM binding strength and its wildtype cognate bound to the same allele with greater than 500 nM binding strength.

### Tumor Clonal Composition Analysis

For the clonal evolution analysis, somatic single-nucleotide variants (SNVs) were called using Mutect 1.1.7^64^ and the variant assurance pipeline (VAP)^66^ for filtering and rescuing. The VAP filters for FFPE and other artifacts and also leverages sequencing data from related samples to salvage false-negatives that would otherwise occur due to limits of the variant caller. When comparing related sample to study clonal evolution, it is especially important to identify shared mutations including those are present at low frequency in some of the samples. VAFs (variant allele frequencies) were calculated for the detected and rescued variants by dividing the number of reads carrying the variant by the total number of reads spanning that position. For each case, mutations covered by less than 20 reads in any sample were removed, as were mutations where the alternate allele was not supported by at least four reads in at least one sample. TitanCNA^67^ was utilized to define local copy number and purity of the tumor samples. Observed VAFs were adjusted for local copy number and purity using the CHAT^68^ framework in order to generate CCF (cancer cell fraction) estimates for each mutation in each sample. Case su002 was excluded from further clonal evolution analysis because it had a purity of < 15% (as inferred by TitanCNA) in both the pre-treatment and post-treatment samples, reducing the accuracy of imputed CCF values. Next, we used PyClone^69^ to define mutational clusters and assess changes in cluster frequencies across treatment. PyClone’s Dirichlet process clustering was carried out on the functional mutations identified in each case. For case su006, since fewer functional mutations were identified compared to the other cases, all filtered mutations (i.e. including synonymous SNVs) that passed the depth of coverage thresholds described above were used to define mutational clusters. The pyclone beta binomial model was run using default parameters for 10,000 iterations with a burn-in of 1000. For visualization of each case, we plotted PyClone clusters comprising at least 1% of the total number of utilized mutations.

### Data Processing of single-cell RNA-seq libraries

Single-cell RNA-seq reads were aligned to the GRCh38 reference genome and quantified using cellranger count (10X Genomics, version 2.1.0). Filtered gene-barcodes matrices containing only barcodes with UMI counts passing threshold for cell detection were used for further analysis. On average, we obtained reads from 1,862 genes per cell (median: 1,716) and 6,304 unique transcripts per cell (median: 4,777), which is comparable to prior droplet based scRNA-seq studies of human cancers.

### Principal component analysis (PCA) and UMAP clustering

All additional analysis was performed using Seurat (version 2.3.4)^18^. Cells with less than 200 genes detected or greater than 10% mitochondrial RNA content were excluded from analysis, with 79,046/83,583 cells passing filter (95%).

For clustering of all cell types in BCC TME, raw UMI counts were log normalized and variable genes called on each dataset independently based on average expression greater than 0.1 and average dispersion greater than 1. Variable T cell receptor and immunoglobulin genes were removed from the list of variable genes to prevent clustering based on variable V(D)J transcripts. To remove batch effects between samples associated with a heat shock gene expression signature, we assigned a heat shock score using the AddModuleScore function based on genes annotated with the GO biological process ontology term ‘cellular response to heat’. Additionally, we assigned scores for S and G2/M cell cycle phase based on previously defined gene sets^24^ using the CellCycleScoring function. Scaled z-scores for each gene were calculated using the ScaleData function and regressed against number of UMIs per cell, mitochondrial RNA content, S phase score, G2/M phase score and heat shock score. Scaled data was used an input into PCA based on variable genes. Clusters were identified using shared nearest neighbor (SNN) based clustering based on the first 20 PCs with k = 30 and resolution = 0.4. The same principal components were used to generate the UMAP projections^19, 20^, which were generated with a minimum distance of 1 and 20 neighbors.

For malignant cell and T cell specific clustering in BCC samples, we isolated subsets of cells from the complete data set identified as either malignant or T cells based on broad clustering. Cells were then re-clustered as described above, with the following modifications: For malignant cells, we did not observe cell-cycle associated effects and did not regress out cell cycle scores. Variable genes were called on the merged dataset based on average expression greater than 0.1 and average dispersion greater than 1.8. For UMAP visualization, we used the first 10 PCs, a minimum distance of 0.15 and 15 neighbors. For T cell clustering, we called variable genes on each dataset independently based on average expression greater than 0.15 and average dispersion greater than 3, and used the merged variable gene set for PCA. T cell clusters were identified using SNN-based clustering based on the first 16 PCs with k = 30 and resolution = 0.3. For UMAP visualization, we used the same PCs, a minimum distance of 0.05 and 20 neighbors.

T cell clustering of SCC samples was performed as described above with the following modifications: Variable genes were called on each dataset independently based on average expression greater than 0.15 and average dispersion greater than 2 and the merged variable gene set used for PCA. We observed three small outlier clusters based on initial clustering which expressed B cell and macrophage marker genes which were removed from further analysis. T cell clusters were identified using SNN-based clustering based on the first 16 PCs with k = 30 and resolution = 0.3. For UMAP visualization, we used the first 16 PCs, a minimum distance of 0.05 and 20 neighbors.

### Cell Cluster Annotation

Clusters were annotated based on expression of known marker genes, including *CD3G, CD3D, CD3E, CD2* (T cells), *CD8A, GZMA* (CD8^+^ T cells), *CD4, FOXP3* (CD4^+^ T cells/Tregs), *KLRC1, KLRC3* (NK cells), *CD19, CD79A* (B cells), *SLAMF7, IGKC* (Plasma cells), *FCGR2A, CSF1R* (Macrophages), *FLT3* (Dendritic cells), *CLEC4C* (pDCs), *COL1A2* (Fibroblasts), *MCAM, MYLK* (Myofibroblasts), *FAP, PDPN* (CAFs), *EPCAM, TP63* (Malignant cells), *PECAM1, VWF* (Endothelial cells), *PMEL, MLANA* (Melanocytes). Clusters were also confirmed by identifying differentially expressed marker genes for each cluster and comparing to known cell type marker genes. Finally, we downloaded bulk RNA-seq count data from sorted immune cell populations from Calderon, *et al*.^21^ and compared bulk gene expression to pseudo-bulk expression profiles from single cell clusters. UMI counts were summed for all cells in each cluster to generate pseudo-bulk profiles. Gene counts from aggregated single-cell and bulk data were then normalized and depth corrected using variance stabilizing transformation in DESeq2 (version 1.18.1). Genes with a coefficient of variation greater than 20% across bulk RNA-seq datasets were used to calculate the Pearson correlation between bulk datasets and pseudo-bulk profiles.

### Data Processing of single-cell TCR-seq libraries

TCR reads were aligned to the GRCh38 reference genome and consensus TCR annotation was performed using cellranger vdj (10X Genomics, version 2.1.0). TCR libraries were sequenced to a minimum depth of 5,000 reads per cell, with a final average of 15,341 reads per cell. On average, 12,335 reads mapped to either the *TRA* or *TRB* loci in each cell. TCR annotation was performed using the 10X cellranger vdj pipeline as described. 85% of annotated T cells were assigned a TCR and only 0.18% of cells not annotated as T cells were assigned a TCR. For cells with two confident TCRs, both were considered in the analysis. Overall, 5.5% of T cells with TCR reads were assigned two productive *TRB* sequences and 5.7% of T cells with TCR reads were assigned two productive TRA sequences and both sequences were assigned to each cell and used for clonotype grouping. Only 1.6% of all cells were assigned two *TRB* sequences and two *TRA* sequences. We detected an average of 1,863 unique clonotypes on average in each patient (range 151 – 4,081). Of 27,956 total clonotypes detected, an average of 1.84 cells were assigned to each clonotype, 5,291 clonotypes comprised of greater than one cell, and clonotype sizes ranging from 1 cell to 564 cells.

### GLIPH Analysis

To identify TCR specificity groups, GLIPH analysis was carried out as described previously^44^. GLIPH clusters TCRs based on two similarity indexes: 1) global similarity, meaning that CDR3 sequences differ by up to one amino acid, and 2) local similarity, meaning that two TCRs contain a common CDR3 motif of 2, 3, or 4 amino acids (enriched over random sub-sampling of unselected repertoires). We performed GLIPH with the following modifications: 1) for clusters based on global similarity, CDR3b fragments within the same cluster are required to be at most one amino acid different, and this difference must be at the same amino acid location in all fragments within the cluster, and 2) for clusters based on local motifs, the starting positions of motifs of the same cluster within CDR3b fragments must be within 3 amino acids to be considered.

### Single-cell CNV detection

Single-cell CNVs were detected using HoneyBADGER^22^. Log-transformed UMI counts were used as input, after removing genes with mean expression lower than 0.1 normalized counts (7,189 genes passing filter, 75-753 genes per chromosome). Non-immune, non-malignant cells were used as a normal reference, including fibroblasts, endothelial cells and melanocytes (n = 2,122). CNVs were detected based on the average gene expression in sliding windows across each chromosome (*n* = 101 genes/window) relative to average expression in normal reference cells. CNV profiles of malignant and reference cells were visualized with z-score limits of -0.6 and 0.6.

### Generation and Data Processing of bulk RNA-seq libraries

For bulk CD4^+^ T cell subset RNA-seq, cDNA library construction was performed using the SMART-Seq v4 Ultra Low Input RNA Kit (Clontech) with 2ng of input RNA. Sequencing libraries were prepared using the Nextera XT DNA Library Prep Kit (Illumina), quantitated using the Qubit dsDNA HS Kit (Thermo Fisher Scientific), and pooled in equimolar ratios. Final pooled libraries were sequenced on an Illumina HiSeq 2500 with paired-end 50-bp read lengths. Paired end RNA-seq libraries from basal cell carcinoma tumors (Atwood *et al*., 2015^30^), cutaneous squamous cell carcinoma tumors (Hoang, *et al*., 2017^31^) and T cell subsets (Simoni, *et al*. 2018^13^ and this study) were aligned to the GRCh38 reference genome using STAR (version 2.6.1a) following adapter trimming by cutadapt (version 1.17). Uniquely-mapped reads were counted using featureCounts (version 1.6.2) using Ensembl GRCh38 GTF transcript annotations. Differential expression analysis was performed using DESeq2 to identify cell-type specific expression programs (version 1.18.1).

### Gene expression signature scoring

Individual cells were scored for bulk RNA-seq expression programs derived from bulk RNA-seq data as follows. Raw UMI counts were used as input into the AUCell package to score each cell for gene set enrichment based on AUC scores to correct for gene dropout and library size differences^70^. After building a gene expression ranking for each cell, the gene set enrichment was calculated for each cell using the area under the recovery curve using default parameters.

Activation and exhaustion signatures were derived by identifying variable genes across all CD8^+^ T cells using the FindVariableGenes function in Seurat with an average expression cutoff of 0.05 and dispersion cutoff of 0.5. The Pearson correlation between reference genes *IFNG* (activation signature) and *HAVCR2* (exhaustion signature) and all variable genes across all CD8^+^ T cells was computed using scaled expression values. Exhaustion and activation signature genes were comprised of the top 50 genes with the highest correlation with reference genes *IFNG* and *HAVCR2*. The *TCF7*^+^/stem-like signature was obtained from processed data from Im *et al*., 2016^45^. Individual cells were scored for enrichment of gene signatures using the function AddModuleScore in Seurat. Cell cycle scoring was performed as previously described^24^. Briefly, cells were scored for enrichment of cell cycle associated genes using the CellCycleScoring function in Seurat.

### Diffusion map and pseudotime analysis

Single cells from BCC samples assigned to CD8^+^ T cell clusters were used for diffusion map and pseudotime analysis. Differentially expressed genes were used to recalculate principal components. Data was then exported to Scanpy (version 1.2.2)^71^ for diffusion map and pseudotime analysis. Data was preprocessed by computing a neighborhood graph using 40 neighbors, the first 20 PCs. The first three components of the diffusion map were then computed. A randomly selected naïve T cell was used as the root cell for diffusion psuedotime computation using the first 3 diffusion components and a minimum group size of 10.

### Sources for bulk RNA sequencing data

Reference bulk RNA-seq from sorted immune populations were obtained from GEO (GSE118165). Reference bulk RNA-seq data from CD8^+^ T cells were obtained from GEO (GSE113590). Reference bulk RNA-seq data from basal cell carcinomas were obtained from GEO (GSE58377). Reference bulk RNA-seq data from squamous cell carcinomas were obtained from the ArrayExpress database (E-MTAB-5678).

### Data Availability

All ensemble and single-cell RNA sequencing data have been deposited in the Gene Expression Omnibus (GEO) under accession GSE123814. Exome sequencing data has been deposited in the Sequence Read Archive (SRA) under accession PRJNA533341.

---

**Supplementary Figure 1. Mutational landscape of BCC tumors following PD-1 blockade, Related to Figure 1. (a)** Summary of mutation burden, potential driver mutations, and mutation frequencies detected in WES data. Potential driver mutations were selected based on frequently mutated genes in BCC identified by Bonilla, *et al*.^16^. **(b)** Bar plots of nonsynonymous mutation burden pre- and post-treatment detected by exome sequencing (top) and predicted neoepitope burden using only the predicted binding strength of the mutant peptide, for peptides with <500 nM binding strength (left), or <50 nM binding strength (right). **(c)** Changes in clonal mutation composition detected in exome sequencing data following treatment, with persistent mutation clusters in grey, mutation clusters decreasing in cellular prevalence following treatment in blue or green, and mutation clusters increasing in cellular prevalence following treatment in red. For clonal composition analysis, variant allele information from matched pre- and posttreatment tumor samples was leveraged to rescue shared low-frequency variants that did not pass standard variant filtering (details in Methods). Bar plots of the ratio of predicted neoepitopes to nonsynomymous mutations in each mutation cluster (right). Predicted neoepitopes were based on binding strength of <500 nM binding strength for the mutant peptide and >500 nM binding strength of the corresponding WT peptide (as in Fig. 1c). **(d)** Representative flow cytometry staining of dissociated BCC cells. Cells were stained for expression of the indicated markers, and two-color histograms are shown for cells pregated as indicated by the arrows and above each diagram. Numbers represent the percentage of cells within the indicated gate. Bottom panels demonstrate cell size differences between malignant and stromal cells, immune cells (non-T cells), and T cells.

**Supplementary Figure 2. Characterization of cell types present in BCC TME, Related to Figure 1. (a)** Heatmap of differentially expressed genes (rows) between cells belonging to each cell type cluster (columns). All malignant cells were treated as one cluster. **(b)** Correlation between immune cell type clusters identified in BCC TME and sorted bulk population from Calderon, *et al*.^21^. UMI counts were summed for each BCC cluster to create pseudo-bulk gene expression profiles and Pearson correlation calculated between each bulk population based on variable genes from bulk profiles. **(c)** UMAP of all BCC TME cells colored by cell type-specific markers. **(d)** Bar plots indicating relative proportions of sort markers detected in each cluster (excluding cells that were not sorted on any markers), relative proportions of cells for which a TCR sequence was detected in each cluster, relative proportions of each non-malignant cell type detected per patient, relative proportions of cells from each patient detected in each cluster, and relative proportions of pre- and post-treatment cells detected in each cluster.

**Supplementary Figure 3. Copy number alterations and gene expression of individual BCC tumors, Related to Figure 1. (a)** Inferred CNV profiles separated by patient based on scRNA-seq (scCNV) and WES. Dashed line indicates a potential subclone identified by scCNV highlighted for su005. For all patients, pre and posttreatment tumor cells were analyzed together and exhibited similar CNV profiles, with the exception of su006. For su006, differences between timepoints were apparent in CNV profiles obtained from both scRNA-seq as well as exome, analogous to the changes in mutation composition identified in Supplementary Figure 1a. **(b)** Heatmap of differentially expressed genes (rows) across malignant BCC cells aggregated by patient (columns). Cutoffs for differential expression were less than 0.01 adjusted p value (Bonferroni corrected), greater than 0.3 average log fold change and greater than 0.3 difference in fraction of positive cells. Core BCC genes that are differentially expressed between all malignant cells and other TME cells are shown in top cluster. Genes differentially expressed between patients are shown in the bottom clusters. Specific genes associated with cancer-associated pathways are highlighted.

**Supplementary Figure 4. Characterization of T cell subtypes present in BCC TME, Related to Figure 2. (a)** Enrichment of bulk T cell subtype signatures for each T cell cluster identified in the BCC TME. T cell subtype signatures were derived from bulk datasets (from this study and Simoni, *et al*.^13^) and single T cells from BCC dataset were scored for signature enrichment. Heatmaps represents the z-scored average signature enrichment for each cluster. **(b)** Heatmap of Pearson correlation between T cell clusters based on first 20 PCs used for clustering. **(c)** UMAP of all T cells colored by subtype-specific markers. Blue indicates high expression and grey indicates low expression. **(d)** UMAP of all T cells separated by patient and colored by anti-PD-1 treatment status.

**Supplementary Figure 5. Characterization of activation/exhaustion trajectories using diffusion maps, Related to Figure 2. (a)** Violin plots of cell coordinates in diffusion components 1 and 2 separated by cluster identity (left, middle). Violin plot of pseudotime values separated by cluster identity (right). **(b)** Heatmap of expression of top correlated genes with diffusion components 1 and 2 (rows) across cells belonging to each cell type cluster (columns).

**Supplementary Figure 6. Increase in Tfh cell clonality following PD-1 blockade accompanied by B cell expansion, Related to Figure 3. (a)** Boxplot of Gini indices for each CD4^+^ T cell cluster separated by timepoint, showing clonal expansion within Tfh cells following treatment. Each point represents a patient with more than 10 cells belonging to a cluster at that timepoint, with the point size proportional to the number of cells. **(b)** UMAP of all cells detected for patient su001 colored by treatment timepoint (left) and relative proportions of each immune cell type (right), showing expansion of B cells post-treatment. **(c)** UMAP of T cells detected for patient su001 colored by treatment timepoint (left) and relative proportions of CD4^+^ phenotype (right), showing expansion of Tfh cells post-treatment. **(d)** Bar plot of percent *AICDA* positive B cells, separated by patient. **(e)** H&E staining of BCC samples from patient su001 post-treatment demonstrating islands of BCC in sclerotic stroma with a peripheral cuff of dense lymphoid tissue. Scale bar for top image represents 400 μm and scale bar for bottom image represents 100 μm.

**Supplementary Figure 7. Correlations between T cell clones or TCR specificity groups and scRNA-seq phenotype, Related to Figure 3. (a)** Distributions of the proportion of cells within each clone (>=3 cells) that share a common cluster identity, separated by patient (for patients with >3 clones with >= 3 cells), compared to randomly selected and size matched groups of T cells (unpaired t-test, two-tailed, \*\*\*\**p*<0.0001, \*\*\**p*<0.001). **(b)** Distribution of the proportion of CD4^+^ cells (left) and CD8^+^ cells (right) within each clone or TCRβ clones within each TCR specificity group (>=3 cells) that share a common cluster identity, separated by treatment timepoint, compared to randomly selected and size matched groups of T cells from the same sample (left, unpaired t-test, two-tailed, \*\*\*\**p*<0.0001, \*\*\**p*<0.001, \*\**p*<0.01, \**p*<0.05). **(c)** Bar plot of T cell cluster assignments for all clones with greater than 10 cells, separated by patient and treatment status. **(d)** Bar plot of T cell cluster assignments for the largest 15 TCR specificity groups, separated by TCRβ clone. Conserved motifs between TCRβ clones identified by GLIPH highlighted in red. Representative TCRβ sequences shown for TCR specificity groups with more than four unique clonotypes. **(e)** Heatmap of the fraction of TCR specificity groups with clones belonging to a given primary phenotype (rows) that also contain clones belonging to a secondary phenotype (columns).

**Supplementary Figure 8. Details of clone transitions, Related to Figure 3. (a)** Heatmap of TCRβ clonotype overlap between all samples, indicating correct pairing of samples and a significant number of overlapping clones between timepoint within individual patients. **(b)** Bar plot of T cell cluster assignments for matched TCRβ clones between timepoints for top 60 clones with at least 3 cells per timepoint. Related to Figure 3e.

**Supplementary Figure 9. Clonal expansion in tumor and peripheral blood detected by bulk TCR sequencing, Related to Figure 4. (a)** Scatterplots comparing TCRβ clone frequencies pre- and post-treatment measured by single-cell RNA_+_TCR-seq, separated by patient. Clones where the majority of cells share an exhausted CD8^+^ phenotype (red) or a memory CD8^+^ phenotype (blue) are highlighted. Patient su003 without no clonotype overlap between timepoints excluded. **(b)** Boxplot of the fraction of novel TCR specificity groups within each cluster following treatment for TCR specificity groups containing at least two distinct TCRβ sequences and at least 3 cells. **(c)** Bar plot of fraction of clones with significant expansion post-treatment based on bulk TCR-seq, separated by patient and phenotype and colored by replacement status. **(d)** Scatterplots comparing TCRβ clone frequencies between timepoints measured by bulk TCR-seq for sequential timepoints in patient su001, with clones where the majority of cells share an exhausted CD8^+^ phenotype (red) or a memory CD8^+^ phenotype (blue) highlighted. Novel clones emerging between timepoints are highlighted in dark red and are detected only in pre- and post-treatment comparisons, but not in comparisons between pre-treatment timepoints, suggesting that replacement is primarily a result of PD-1 blockade rather than time between sampling.

**Supplementary Figure 10. TCR overlap between peripheral blood and tumor detected by bulk TCR sequencing, Related to Figure 4. (a)** Pie chart of percentage of TCRβ clones detected in peripheral blood that were also detected by scRNA-seq, expanded to show distribution of phenotypes in tumor, as well as fraction of exhausted clones detected in peripheral blood, colored by replacement status in tumor. **(b)** Bar plot of percentage peripheral T cells matching tumor-infiltrating TCRβ clones with exhausted phenotypes post-treatment as detected by scRNA-seq. (c) Violin plot of TCR specificity group enrichment (tumor frequency / PBMC frequency) detected by bulk TCR-seq, separated by phenotype and treatment status (unpaired t-test, one-tailed, \*\*\*\**p*<0.0001, \*\*\**p*<0.001, \*\**p*<0.01, \**p*<0.05).

**Supplementary Figure 11. Clonal replacement analysis in SCC TILs following PD-1 blockade, Related to Figure 4. (a)** UMAP of tumor-infiltrating T cells present in SCC samples pre- and post-PD-1 blockade colored by patient (top right) and anti-PD-1 treatment status (bottom right). **(b)** Heatmap of correlation between averaged RNA expression between BCC and SCC T cell clusters. **(c)** Boxplot of Gini indices for each CD8^+^ T cell cluster calculated for each patient. **(d)** Abundance of the top 12 exhausted clones in sample su010-S compared to the abundance of the same clones in sorted CD8^+^ CD39^+^ T cells, colored by assigned phenotype. **(e)** Distribution of the proportion of cells within each clone or TCRβ clones within each TCR specificity group (>=3 cells) that share a common cluster identity, separated by treatment timepoint, compared to randomly selected and size matched groups of T cells from the same sample (left, unpaired t-test, two-tailed, \*\*\*\**p*<0.0001, \*\**p*<0.01, \**p*<0.05). **(f)** Heatmap of the fraction of clonotypes belonging to a given primary phenotype cluster (rows) that are shared with other secondary phenotype clusters (columns). **(g)** Heatmap of all observed phenotype transitions for matched clones during PD-1 blockade for clones with at least 3 cells for each timepoint. **(h)** *TCF7*^+^/stem-like score (from Im *et al*^45^) versus exhaustion score for all CD8^+^ T cells, colored by gene expression (left). *TCF7*^+^/stem-like score versus exhaustion score for exhausted cells and cells of other phenotypes belonging to primarily exhausted clones, colored by phenotype (top right). Violin plot of *TCF7*^+^/stem-like score for exhausted cells and cells of other phenotypes belonging to primarily exhausted clones, demonstrating that the highest *TCF7*^+^/stem-like score is observed in cells with an exhausted phenotype (bottom right). **(i)** Violin plot of *TCF7*^+^/stem-like score for memory and exhausted cells separated by change in clone abundance following treatment (left). **(j)** Scatterplots comparing TCRβ clone frequencies pre- and post-treatment measured by single-cell TCR sequencing for all BCC patients. Clones that were significantly expanded or contracted post-treatment based on a Fisher exact test are highlighted on the left. Clones where the majority of cells share an exhausted CD8^+^ phenotype (middle, red) or a memory CD8^+^ phenotype (right, blue) are also highlighted.

## References

1. Sharma, P. & Allison, J. P. The future of immune checkpoint therapy. Science 348, 56–61 (2015).

2. Page, D. B., Postow, M. A., Callahan, M. K., Allison, J. P. & Wolchok, J. D. Immune modulation in cancer with antibodies. Annu. Rev. Med. 65, 185–202 (2014).

3. Shin, D. S. & Ribas, A. The evolution of checkpoint blockade as a cancer therapy: what’s here, what’s next? Curr. Opin. Immunol. 33, 23–35 (2015).

4. Hamid, O. et al. Safety and tumor responses with lambrolizumab (anti-PD-1) in melanoma. N. Engl. J. Med. 369, 134–144 (2013).

5. Robert, C. et al. Anti-programmed-death-receptor-1 treatment with pembrolizumab in ipilimumab-refractory advanced melanoma: a randomised dose-comparison cohort of a phase 1 trial. Lancet Lond. Engl. 384, 1109–1117 (2014).

6. Wolchok, J. D. et al. Nivolumab plus ipilimumab in advanced melanoma. N. Engl. J. Med. 369, 122–133 (2013).

7. Fourcade, J. etal. Upregulation of Tim-3 and PD-1 expression is associated with tumor antigen-specific CD8+ T cell dysfunction in melanoma patients. J. Exp. Med. 207, 2175–2186 (2010).

8. Sakuishi, K. et al. Targeting Tim-3 and PD-1 pathways to reverse T cell exhaustion and restore anti-tumor immunity. J. Exp. Med. 207, 2187–2194 (2010).

9. Wherry, E. J. & Kurachi, M. Molecular and cellular insights into T cell exhaustion. Nat. Rev. Immunol. 15, 486–499 (2015).

10. Singer, M. et al. A Distinct Gene Module for Dysfunction Uncoupled from Activation in Tumor-Infiltrating T Cells. Cell 166, 1500–1511.e9 (2016).

11. Pauken, K. E. et al. Epigenetic stability of exhausted T cells limits durability of reinvigoration by PD-1 blockade. Science 354, 1160–1165 (2016).

12. Sen, D. R. et al. The epigenetic landscape of T cell exhaustion. Science 354, 1165–1169 (2016).

13. Simoni, Y. et al. Bystander CD8 + T cells are abundant and phenotypically distinct in human tumour infiltrates. Nature 557, 575 (2018).

14. Chang, A. L. S. et al. Pembrolizumab for advanced basal cell carcinoma: an investigator-initiated, proof-of-concept study. J. Am. Acad. Dermatol. (2018). doi:10.1016/j.jaad.2018.08.017

15. Eisenhauer, E. A. et al. New response evaluation criteria in solid tumours: revised RECIST guideline (version 1.1). Eur. J. Cancer Oxf. Engl. 1990 45, 228–247 (2009).

16. Bonilla, X. et al. Genomic analysis identifies new drivers and progression pathways in skin basal cell carcinoma. Nat. Genet. 47, 398 (2015).

17. Hundal, J. et al. pVAC-Seq: A genome-guided in silico approach to identifying tumor neoantigens. Genome Med. 8, 11 (2016).

18. Butler, A., Hoffman, P., Smibert, P., Papalexi, E. & Satija, R. Integrating single-cell transcriptomic data across different conditions, technologies, and species. Nat. Biotechnol. 36, 411–420 (2018).

19. Mclnnes, L., Healy, J. & Melville, J. UMAP: Uniform Manifold Approximation and Projection for Dimension Reduction. ArXiv180203426 Cs Stat (2018).

20. Becht, E. et al. Dimensionality reduction for visualizing single-cell data using UMAP. Nat. Biotechnol. 37, 38–44 (2019).

21. Calderon, D. et al. Landscape of stimulation-responsive chromatin across diverse human immune cells. bioRxiv 409722 (2018). doi:10.1101/409722

22. Fan, J. et al. Linking transcriptional and genetic tumor heterogeneity through allele analysis of single-cell RNA-seq data. Genome Res. gr.228080.117 (2018). doi:10.1101/gr.228080.117

23. Rubin, A. I., Chen, E. H. & Ratner, D. Basal-cell carcinoma. N. Engl. J. Med. 353, 2262–2269 (2005).

24. Tirosh, I. et al. Dissecting the multicellular ecosystem of metastatic melanoma by single-cell RNA-seq. Science 352, 189–196 (2016).

25. Puram, S. V. et al. Single-Cell Transcriptomic Analysis of Primary and Metastatic Tumor Ecosystems in Head and Neck Cancer. Cell 171, 1611–1624.e24 (2017).

26. Tellechea, O., Reis, J. P., Domingues, J. C. & Baptista, A. P. Monoclonal antibody Ber EP4 distinguishes basal-cell carcinoma from squamous-cell carcinoma of the skin. Am. J. Dermatopathol. 15, 452–455 (1993).

27. Bircan, S., Candir, O., Kapucoglu, N. & Baspinar, S. The expression of p63 in basal cell carcinomas and association with histological differentiation. J. Cutan. Pathol. 33, 293–298 (2006).

28. Bernemann, T. M., Podda, M., Wolter, M. & Boehncke, W. H. Expression of the basal cell adhesion molecule (B-CAM) in normal and diseased human skin. J. Cutan. Pathol. 27, 108–111 (2000).

29. Ransohoff, K. J., Tang, J. Y. & Sarin, K. Y. Squamous Change in Basal-Cell Carcinoma with Drug Resistance. (2015). doi:10.1056/NEJMc1504261

30. Atwood, S. X. et al. Smoothened variants explain the majority of drug resistance in basal cell carcinoma. Cancer Cell 27, 342–353 (2015).

31. Hoang, V. L. T. et al. RNA-seq reveals more consistent reference genes for gene expression studies in human non-melanoma skin cancers. PeerJ 5, e3631 (2017).

32. Wei, S. C. et al. Distinct Cellular Mechanisms Underlie Anti-CTLA-4 and Anti-PD-1 Checkpoint Blockade. Cell 170, 1120–1133.e17 (2017).

33. Sade-Feldman, M. et al. Defining T Cell States Associated with Response to Checkpoint Immunotherapy in Melanoma. Cell 175, 998–1013.e20 (2018).

34. Haghverdi, L., Buettner, F. & Theis, F. J. Diffusion maps for high-dimensional single-cell analysis of differentiation data. Bioinforma. Oxf. Engl. 31, 2989–2998 (2015).

35. Kumar, B. V. et al. Human Tissue-Resident Memory T Cells Are Defined by Core Transcriptional and Functional Signatures in Lymphoid and Mucosal Sites. Cell Rep. 20, 2921–2934 (2017).

36. Savas, P. et al. Single-cell profiling of breast cancer T cells reveals a tissue-resident memory subset associated with improved prognosis. Nat. Med. 24, 986 (2018).

37. Duhen, T. et al. Co-expression of CD39 and CD103 identifies tumor-reactive CD8 T cells in human solid tumors. Nat. Commun. 9, 2724 (2018).

38. Li, H. et al. Dysfunctional CD8 T Cells Form a Proliferative, Dynamically Regulated Compartment within Human Melanoma. Cell 176, 775–789.e18 (2019).

39. Thommen, D. S. et al. A transcriptionally and functionally distinct PD-1 + CD8 + T cell pool with predictive potential in non-small-cell lung cancer treated with PD-1 blockade. Nat. Med. 24, 994 (2018).

40. Gros, A. et al. Prospective identification of neoantigen-specific lymphocytes in the peripheral blood of melanoma patients. Nat. Med. 22, 433–438 (2016).

41. Azizi, E. et al. Single-Cell Map of Diverse Immune Phenotypes in the Breast Tumor Microenvironment. Cell 174, 1293–1308.e36 (2018).

42. Zhang, L. et al. Lineage tracking reveals dynamic relationships of T cells in colorectal cancer. Nature 1 (2018). doi:10.1038/s41586-018-0694-x

43. Becattini, S. et al. Functional heterogeneity of human memory CD4+ T cell clones primed by pathogens or vaccines. Science 347, 400–406 (2015).

44. Glanville, J. et al. Identifying specificity groups in the T cell receptor repertoire. Nature 547, 94–98 (2017).

45. Im, S. J. et al. Defining CD8+ T cells that provide the proliferative burst after PD-1 therapy. Nature 537, 417–421 (2016).

46. Kurtulus, S. et al. Checkpoint Blockade Immunotherapy Induces Dynamic Changes in PD-1-CD8+ Tumor-Infiltrating T Cells. Immunity 50, 181–194.e6 (2019).

47. Siddiqui, I. et al. Intratumoral Tcf1+PD-1+CD8+ T Cells with Stem-like Properties Promote Tumor Control in Response to Vaccination and Checkpoint Blockade Immunotherapy. Immunity 50, 195–211.e10 (2019).

48. Miller, B. C. et al. Subsets of exhausted CD8 + T cells differentially mediate tumor control and respond to checkpoint blockade. Nat. Immunol. 20, 326 (2019).

49. Huang, A. C. et al. A single dose of neoadjuvant PD-1 blockade predicts clinical outcomes in resectable melanoma. Nat. Med. 25, 454 (2019).

50. Kamphorst, A. O. et al. Proliferation of PD-1+ CD8 T cells in peripheral blood after PD-1-targeted therapy in lung cancer patients. Proc. Natl. Acad. Sci. U. S. A. 114, 4993–4998 (2017).

51. Migden, M. R. et al. PD-1 Blockade with Cemiplimab in Advanced Cutaneous Squamous-Cell Carcinoma. N. Engl. J. Med. 379, 341–351 (2018).

52. Scheper, W. et al. Low and variable tumor reactivity of the intratumoral TCR repertoire in human cancers. Nat. Med. 25, 89–94 (2019).

53. Ghoneim, H. E. et al. De Novo Epigenetic Programs Inhibit PD-1 Blockade-Mediated T Cell Rejuvenation. Cell 170, 142–157.e19 (2017).

54. Spitzer, M. H. et al. Systemic Immunity Is Required for Effective Cancer Immunotherapy. Cell 168, 487–502.e15 (2017).

55. Chamoto, K. et al. Mitochondrial activation chemicals synergize with surface receptor PD-1 blockade for T cell-dependent antitumor activity. Proc. Natl. Acad. Sci. U. S. A. 114, E761–E770 (2017).

56. Li, J. et al. Tumor Cell-Intrinsic Factors Underlie Heterogeneity of Immune Cell Infiltration and Response to Immunotherapy. Immunity 49, 178–193.e7 (2018).

57. Matsushita, H. et al. Cancer exome analysis reveals a T-cell-dependent mechanism of cancer immunoediting. Nature 482, 400–404 (2012).

58. Gee, M. H. et al. Antigen Identification for Orphan T Cell Receptors Expressed on Tumor-Infiltrating Lymphocytes. Cell (2017). doi:10.1016/j.cell.2017.11.043

59. Wang, C. et al. High-throughput, high-fidelity HLA genotyping with deep sequencing. Proc. Natl. Acad. Sci. U. S. A. 109, 8676–8681 (2012).

60. Thorstenson, Y. R. et al. Allelic resolution NGS HLA typing of Class I and Class II loci and haplotypes in Cape Town, South Africa. Hum. Immunol. (2018). doi:10.1016/j.humimm.2018.09.004

61. McKenna, A. et al. The Genome Analysis Toolkit: a MapReduce framework for analyzing next-generation DNA sequencing data. Genome Res. 20, 1297–1303 (2010).

62. Li, H. & Durbin, R. Fast and accurate short read alignment with Burrows-Wheeler transform. Bioinformatics 25, 1754–1760 (2009).

63. Birger, C. et al. FireCloud, a scalable cloud-based platform for collaborative genome analysis: Strategies for reducing and controlling costs. bioRxiv 209494 (2017). doi:10.1101/209494

64. Cibulskis, K. et al. Sensitive detection of somatic point mutations in impure and heterogeneous cancer samples. Nat. Biotechnol. 31, 213–219 (2013).

65. Xie, C. et al. Fast and accurate HLA typing from short-read next-generation sequence data with xHLA. Proc. Natl. Acad. Sci. 114, 8059–8064 (2017).

66. Sun, R. et al. Between-region genetic divergence reflects the mode and tempo of tumor evolution. Nat. Genet. 49, 1015–1024 (2017).

67. Ha, G. et al. TITAN: inference of copy number architectures in clonal cell populations from tumor whole-genome sequence data. Genome Res. 24, 1881–1893 (2014).

68. Li, B. & Li, J. Z. A general framework for analyzing tumor subclonality using SNP array and DNA sequencing data. Genome Biol. 15, 473 (2014).

69. Roth, A. et al. PyClone: statistical inference of clonal population structure in cancer. Nat. Methods 11, 396–398 (2014).

70. Aibar, S. et al. SCENIC: single-cell regulatory network inference and clustering. Nat. Methods 14, 1083–1086 (2017).

71. Wolf, F. A., Angerer, P. & Theis, F. J. SCANPY: large-scale single-cell gene expression data analysis. Genome Biol. 19, 15 (2018).

