## Supplementary Figures for "Clonal replacement of tumor-specific T cells following PD-1 blockade"

#### Supplementary Figure 1

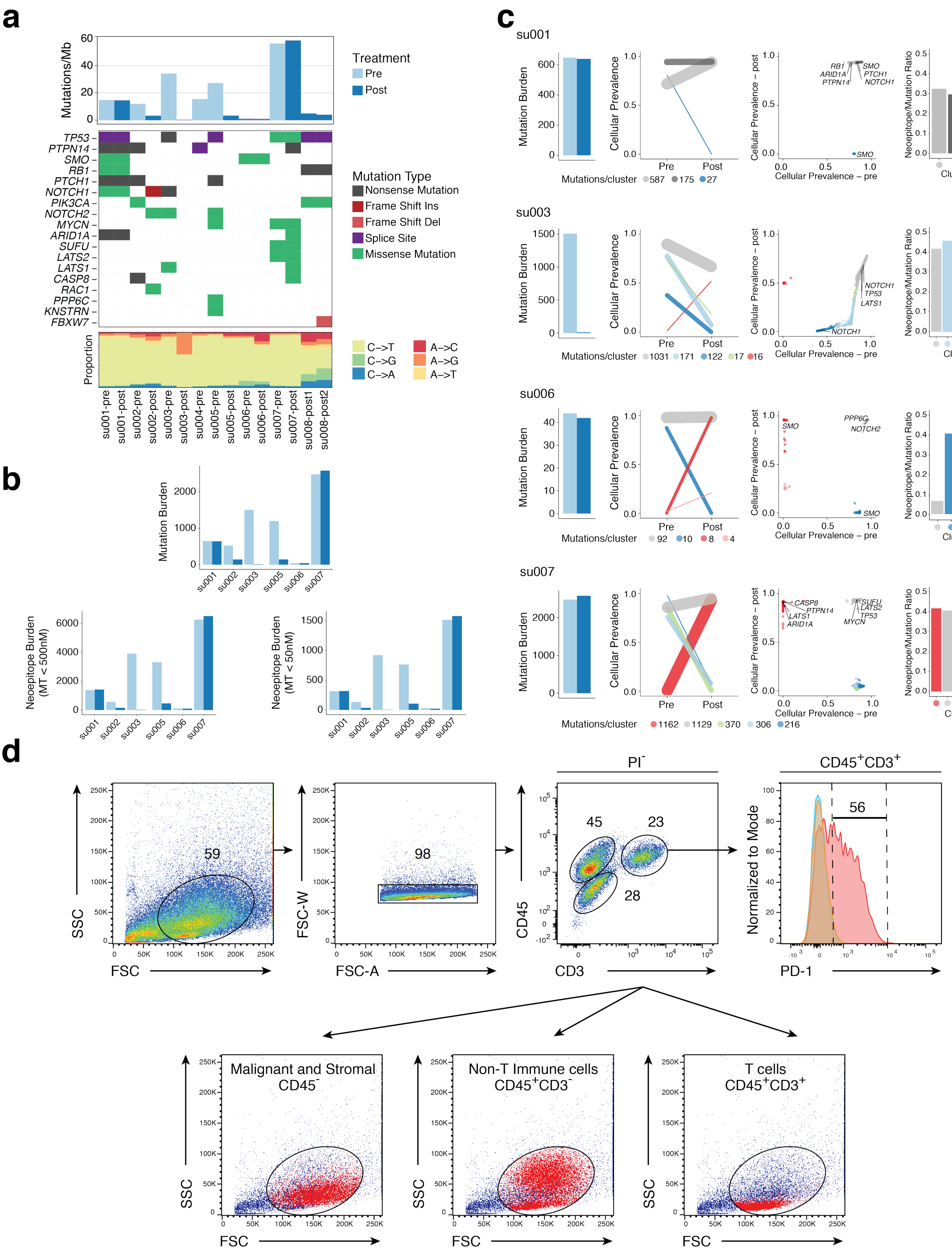

#### Supplementary Figure 2

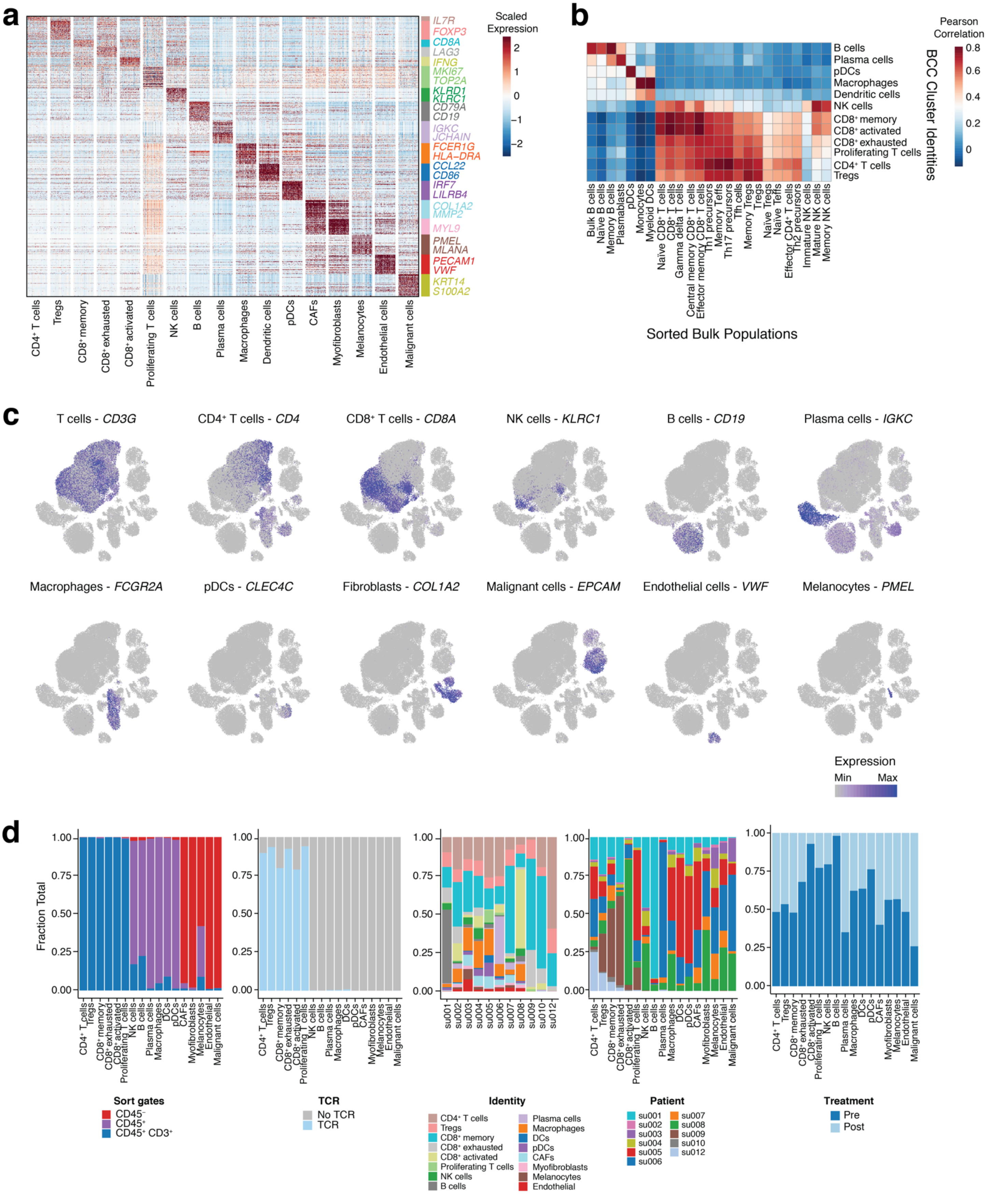

### Supplementary Figure 3

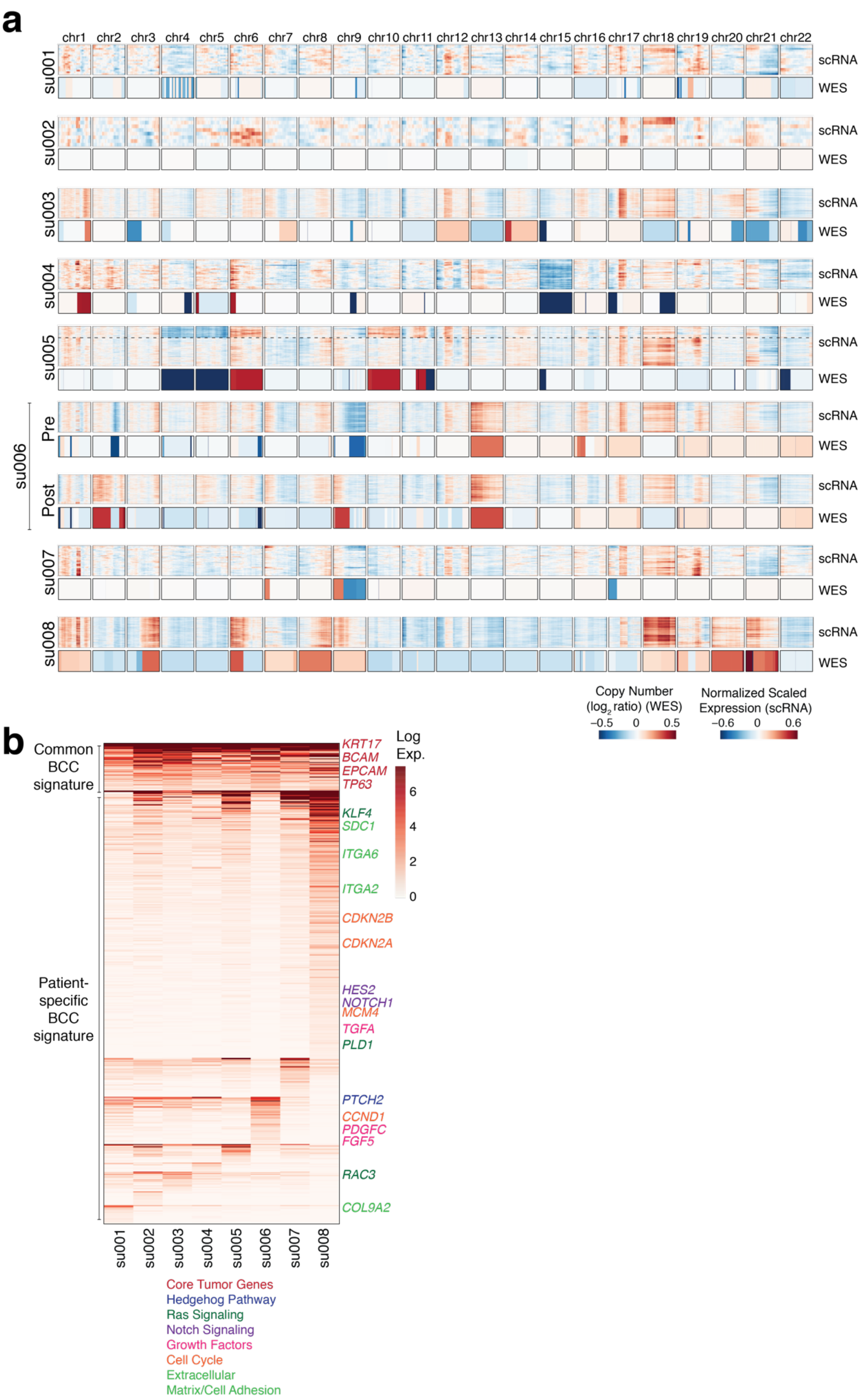

### Supplementary Figure 4

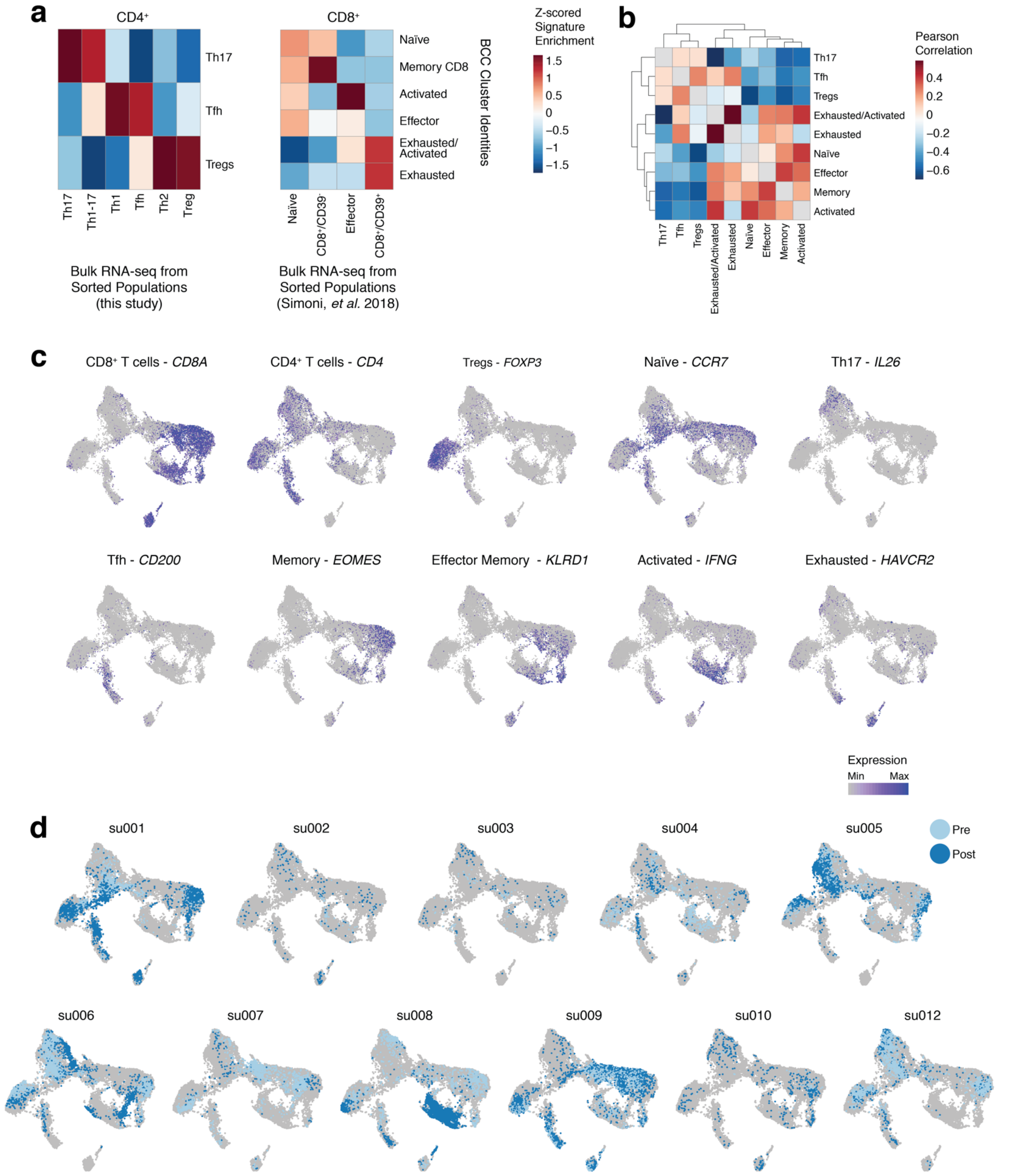

#### Supplementary Figure 5

**a**

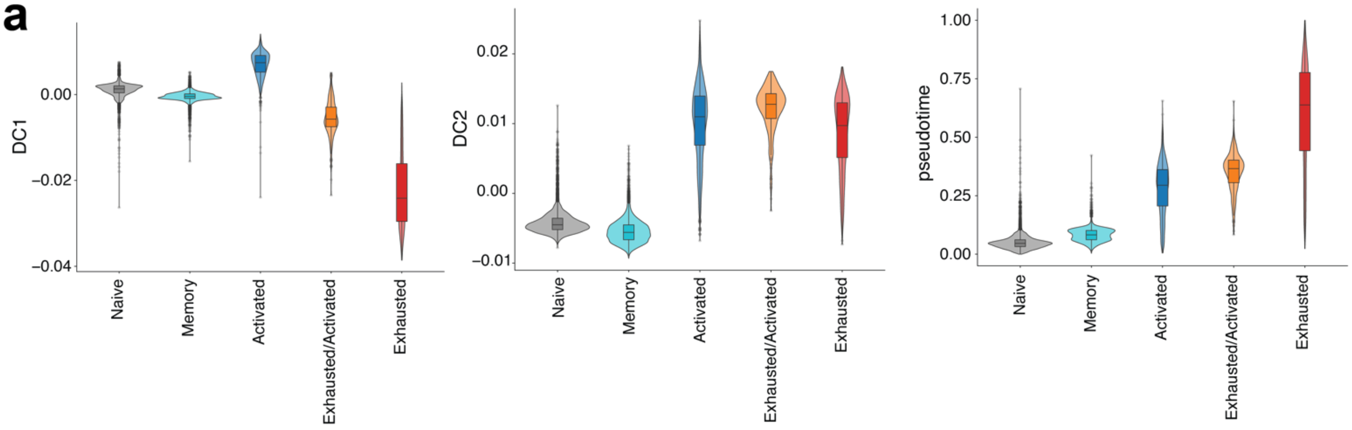


**b**

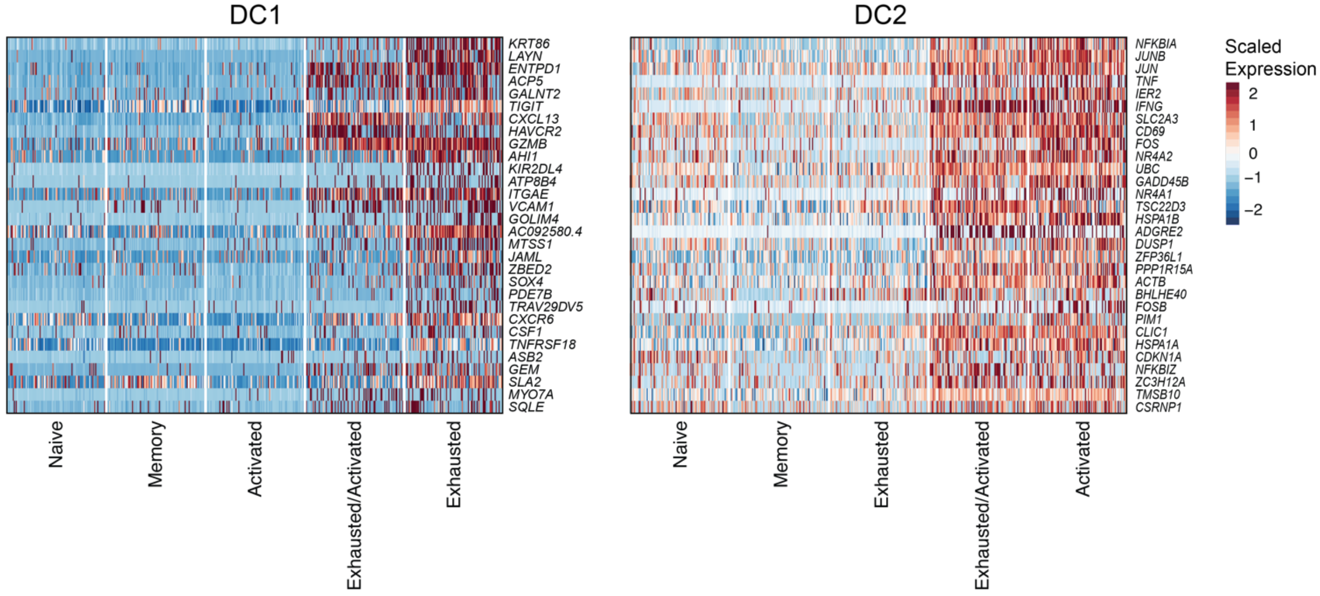

#### Supplementary Figure 6

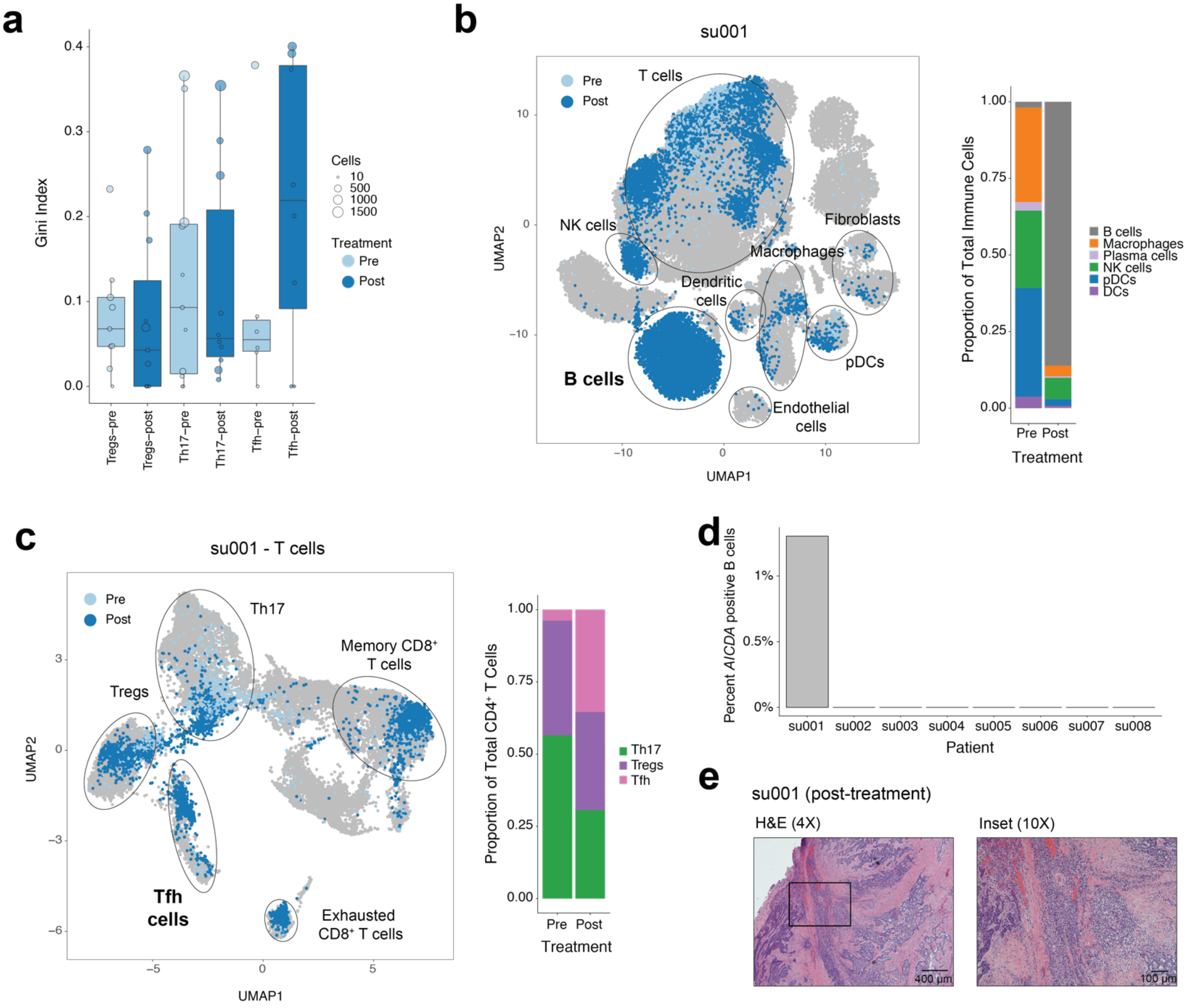

#### Supplementary Figure 7

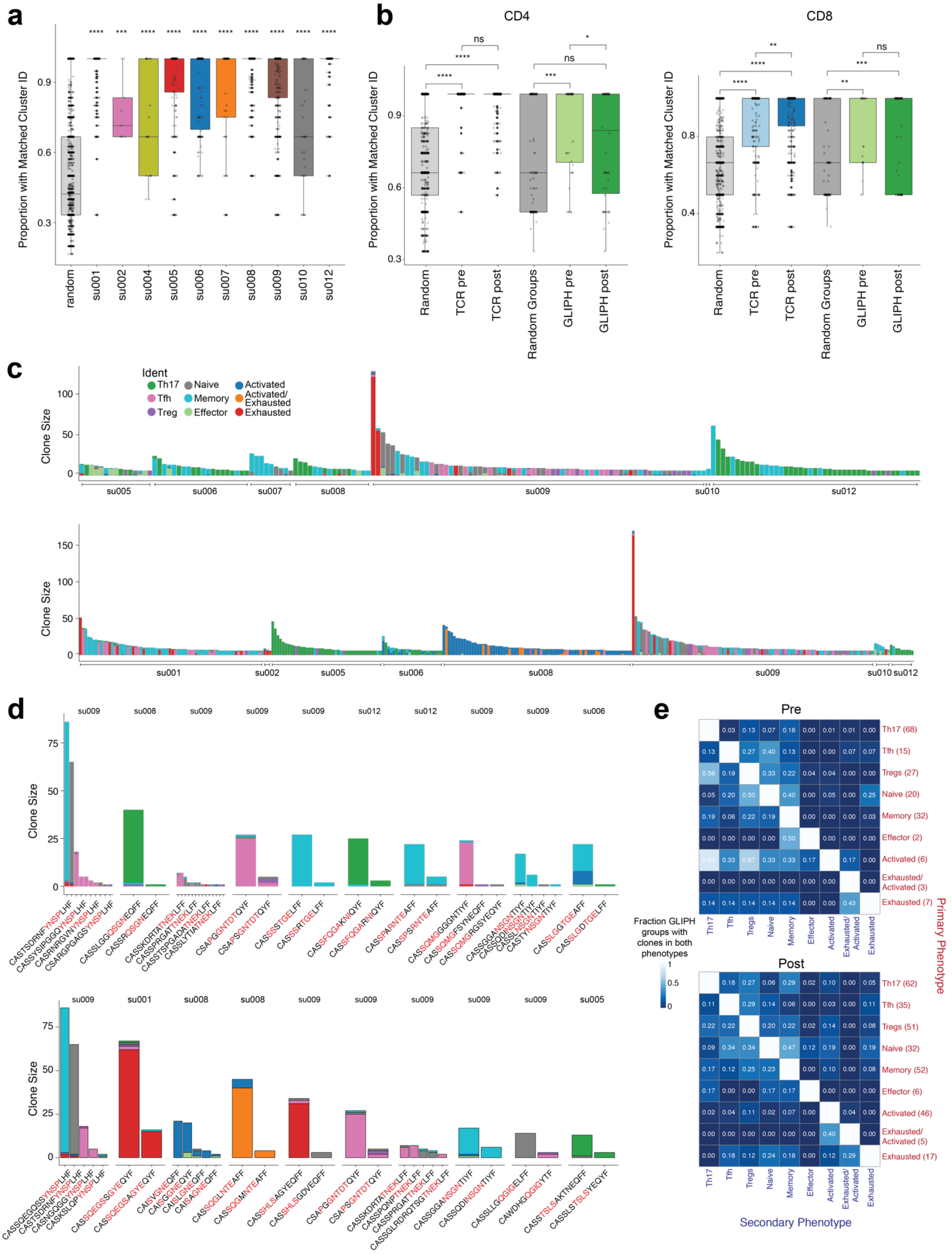

### Supplementary Figure 8

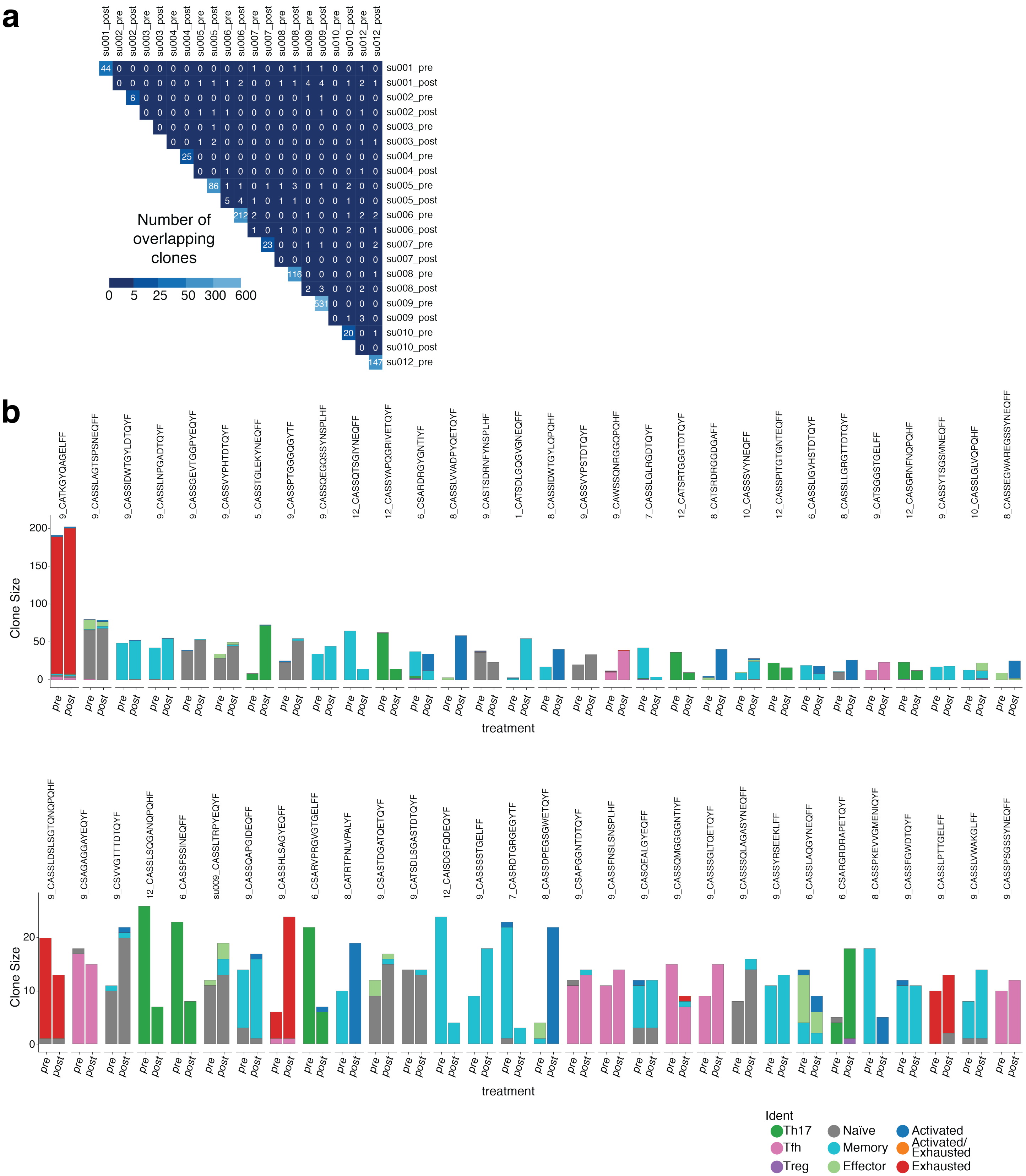

### Supplementary Figure 9

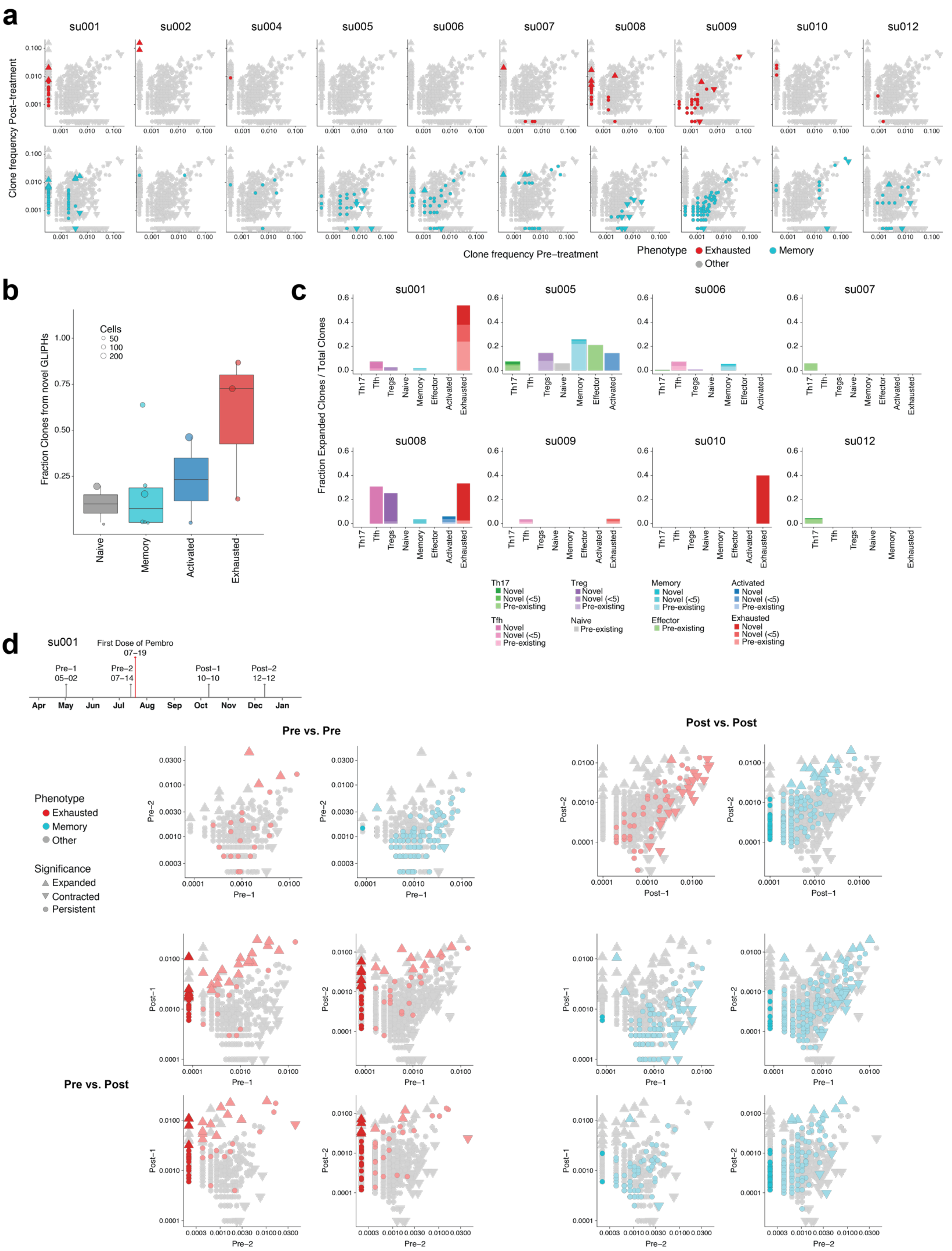

### Supplementary Figure 10

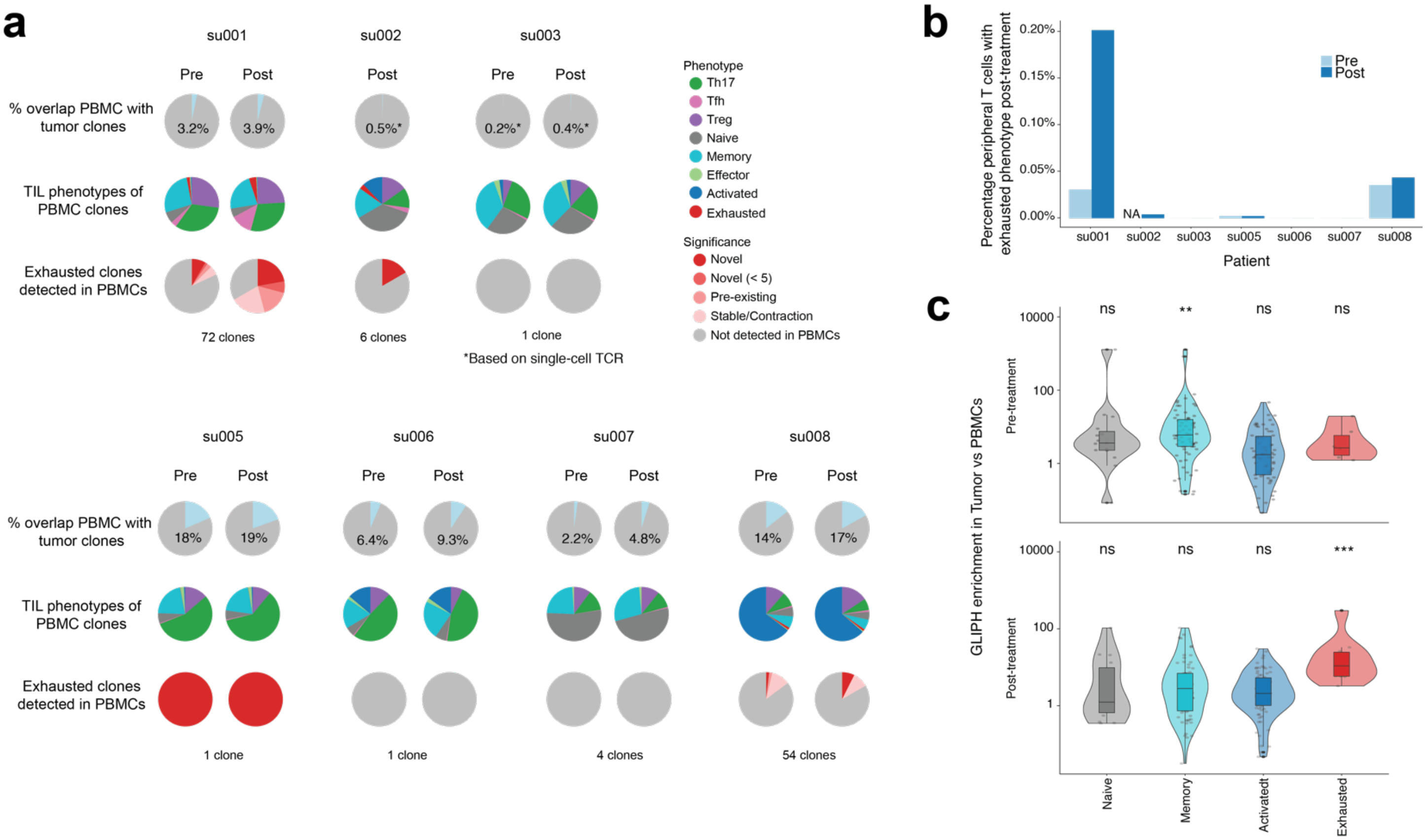

#### Supplementary Figure 11

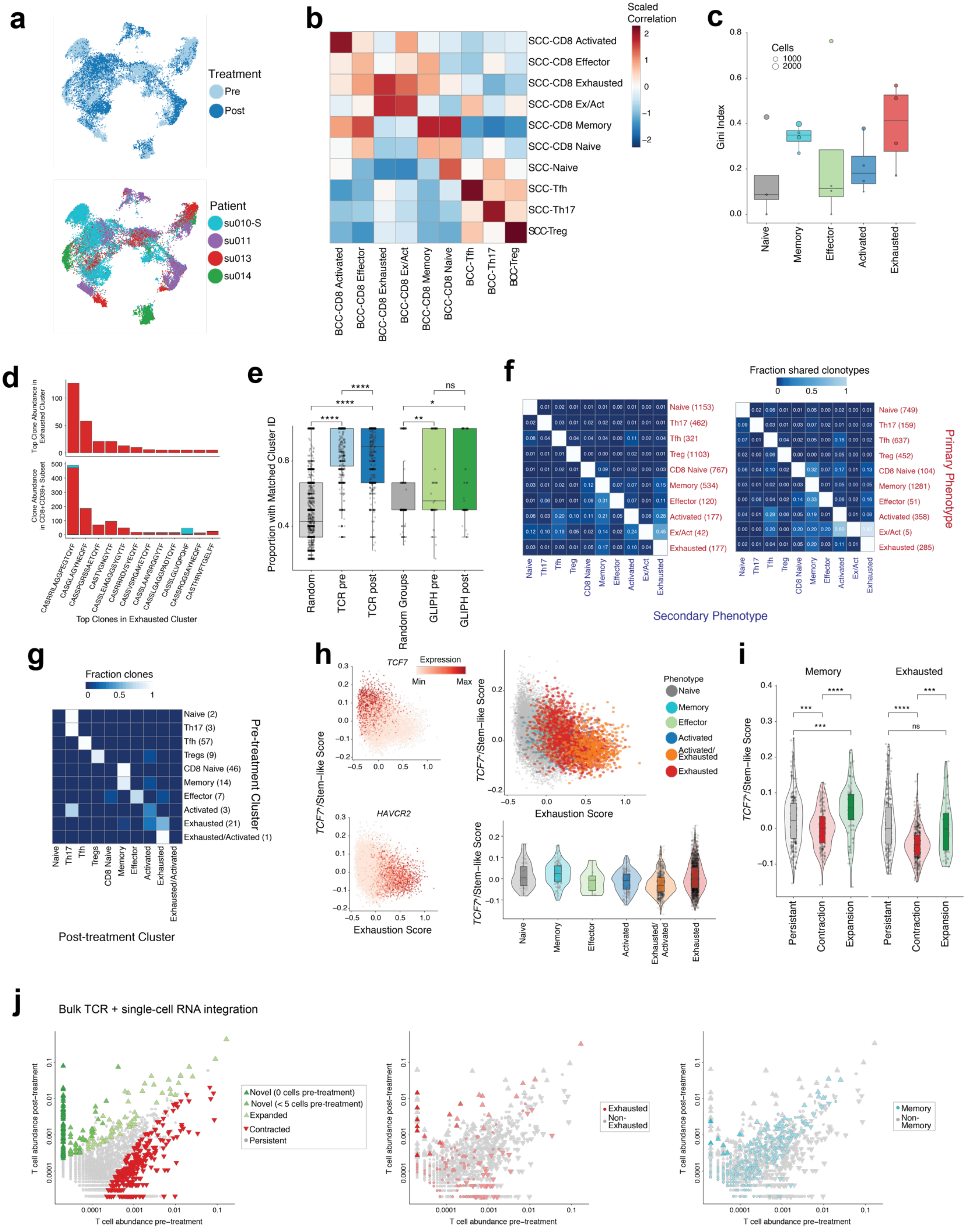
